# Spatiotemporal dynamics of locomotor decisions in *Drosophila melanogaster*

**DOI:** 10.1101/2024.09.04.611038

**Authors:** Lior Lebovich, Tom Alisch, Edward S. Redhead, Matthew O. Parker, Yonatan Loewenstein, Iain D. Couzin, Benjamin L. de Bivort

**Affiliations:** Department of Collective Behaviour, Max Planck Institute of Animal Behavior, Konstanz, Germany; Centre for the Advanced Study of Collective Behaviour, University of Konstanz, Universitätsstraße 10, 78464, Konstanz, Germany; Department of Biology, University of Konstanz, Konstanz, Germany; Department of Organismic & Evolutionary Biology & Center for Brain Science, Harvard University, Cambridge, Massachusetts, U.S.A.; School of Psychology, University of Southampton, UK; School of Biosciences, University of Surrey, UK; The Edmond and Lily Safra Center for Brain Sciences, The Alexander Silberman Institute of Life Sciences, Dept. of Cognitive and Brain Sciences and The Federmann Center for the Study of Rationality, The Hebrew University of Jerusalem, Jerusalem, Israel

**Author notes:** equal contribution (Lior Lebovich, Tom Alisch). equal contribution (Yonatan Loewenstein, Iain D. Couzin). **Email addresses:** LL; TA; ER; MP; YL; IC; BdB. **Author Contributions:** Designed experiments: LL, TA, ER, MP, YL, IC, BdB Performed research: LL, TA, ER Contributed new reagents or analytic tools: ER, MP Analyzed data: LL, TA, ER, MP, YL, IC, BdB Wrote the paper: LL, TA, ER, MP, YL, IC, BdB. **Competing Interest Statement:** The authors declare no competing interests.

**Keywords:** Decision-making, Y-maze, *Drosophila melanogaster*, embodied choice, movement decisions

## Abstract

Decision-making in animals often involves choosing actions while navigating the environment, a process markedly different from static decision paradigms commonly studied in laboratory settings. Even in decision-making assays in which animals can freely locomote, decision outcomes are often interpreted as happening at single points in space and single moments in time, a simplification that potentially glosses over important spatiotemporal dynamics. We investigated locomotor decision-making in *Drosophila melanogaster* in Y-shaped mazes, measuring the extent to which their future choices could be predicted through space and time. We demonstrate that turn-decisions can be reliably predicted from flies’ locomotor dynamics, with distinct predictability phases emerging as flies progress through maze regions. We show that these predictability dynamics are not merely the result of maze geometry or wall-following tendencies, but instead reflect the capacity of flies to move in ways that depend on sustained locomotor signatures, suggesting an active, working memory-like process. Additionally, we demonstrate that fly mutants known to have sensory and information-processing deficits exhibit altered spatial predictability patterns, highlighting the role of visual, mechanosensory, and dopaminergic signaling in locomotor decision-making. Finally, highlighting the broad applicability of our analyses, we generalize our findings to other species and tasks. We show that human participants in a virtual Y-maze exhibited similar decision predictability dynamics as flies. This study advances our understanding of decision-making processes, emphasizing the importance of spatial and temporal dynamics of locomotor behavior in the lead-up to discrete choice outcomes.

## Introduction

Decision-making is the process of choosing an action among a set of possible alternatives. The scientific literature concerning decision-making processes is dominated by paradigms in which an individual decision-maker is stationary and makes a static choice (e.g., Hanks et al., 2015). But in naturalistic conditions, response behavior is typically much more fluid. From foraging to courtship to finding refuge, animals make countless, vital locomotor decisions that can vary continuously in space with little constraint on the space of actions. For example, they might navigate upwind along an encountered odor plume to find a nearby nutrient-rich substrate (Giang et al., 2017), or move to avoid a looming predator (Gibson et al., 2015; Kempraj et al., 2020). Such contexts require decision-makers to not only implement the decision while moving through the environment, but also to process information about the upcoming decision and adjust their choice-behavior and, hence, movement trajectory on the go (or rather, on the fly).

Decision-making experiments can be separated into two categories based on whether the decision-maker is static or on the move. Examples of static decision-making assays come from many species and include, for example, human participants pulling levers in accordance with random-dot movement direction (Selen et al., 2012), tethered mosquitoes steering towards visual stimuli (Vinauger et al., 2019), head-restrained mice detecting odors in an olfactory figure-background segregation task (Lebovich et al., 2021), tethered zebrafish larvae modulating swimming direction in accordance with visual background movement (Bahl & Engert, 2020), macaques making saccadic eye movements in accordance with random-dot motion (Huk & Shadlen, 2005; Shadlen & Kiani, 2013), or head-fixed rats licking water spouts upon detected whisker deflections (Stüttgen & Schwarz, 2010). Assays of decision-making on the move encompass a comparable range of animals and include, for example, auditory-guided decision-making in rats (Jankowski et al., 2023), olfaction-mediated steering behavior of fruit fly larvae (Lin et al., 2019), odor-mediated steering behavior of nematodes (Liu et al., 2018), social isolation-mediated turning behavior of rats (Kim et al., 2023), or visual stimulus-induced optomotor response in fruit flies (Werkhoven et al., 2021).

In lab experimental settings, decision-making on the move is typically assayed in “arenas”, structures that allow free movement but only within static boundaries that ultimately constrain locomotion (but see Sridhar et al., 2021). Arenas have many different general forms and specific geometries, and may incorporate distinct locational features, depending on task requirements (Wijnen et al., 2024). For example, foraging decisions of bumble bees have been probed using a radial arm maze, featuring four symmetric centrally intersecting chambers (Pull et al., 2022), memory-dependent search decisions of mice have been investigated using a cheeseboard maze, a circular arena featuring equally spaced wells (Dupret et al., 2010), and social proximity decisions have been examined in circular arenas with a linear barrier separating two conspecific flies (Alisch et al., 2018). In some decision paradigms, arenas that are relatively large and minimize distinct spatial features are used to approximate behavior in unconstrained settings. For example, nociceptive stimulus-dependent initiation of stereotyped locomotor behavior of *Drosophila melanogaster* larvae (Ohyama et al., 2015), odor-mediated modulation of aversive locomotor behavior of *C. elegans* (Liu et al., 2020), and anxiety-related wall-following thigmotaxis of *M. musculus* (Seibenhener & Wooten, 2015) have been explored in square or circular open field arenas. Across maze types, it is often either explicitly tested (Mai et al., 2012), or implicitly assumed, that organisms move through their environment and make decisions in a goal-directed manner based on motivational states, such as exploration or escape (Thomas et al., 2014).

One form of arena that is particularly useful for studying two-choice decision-making on the move is the so-called Y-maze (Cleal et al., 2020). Its geometry comprises 3 arms, typically of equal length, that meet in a single intersection and are often separated by 120° (see e.g., Warden et al., 1929). This configuration permits repeated readouts of binary locomotor decisions. An organism traversing along a maze arm towards the intersection and then turning into either the left or the right arm constitutes a binary locomotor decision. Y-mazes have historically been utilized to study various forms of decision-making across many organisms, such as effort-dependent foraging decisions in *M. musculus* (Dylda & Wang, 2022), expression of learned color-reward association through turning decisions in the hornet *Vespa crabro* (Lacombrade et al., 2023), visual-cue dependent navigational decisions in the golden shiner fish *Notemigonus crysoleucas* (Woodley et al., 2019), or handedness bias (Ayroles et al., 2015; Buchanan et al., 2015) and associative conditioning (Adel & Griffith, 2021; Lesar et al., 2021) in *D. melanogaster*.

A critical feature of Y-mazes (and other spatial arenas) is the location at which a decision is ultimately recorded as an outcome, typically the point at which the maze arms intersect. The test subject’s decision can be easily labeled based on their behavior moving through this “choice point” and the effect of experimental variables assessed. For instance, prominent behavioral features predicting decision outcomes include the history of presented stimuli (Yeshurun et al., 2008; Ashourian & Loewenstein, 2011; Raviv et al., 2012), past decisions (Urai et al., 2019) and their outcomes (Thorndike, 1911; Skinner, 1938; Ferster & Skinner, 1957; Shteingart & Loewenstein, 2014; Mongillo et al., 2014), as well as individual choice preference (Kain et al., 2012; Buchanan et al., 2015; Gorostiza et al., 2016; de Bivort et al., 2019; Lebovich et al., 2019; Churgin et al., 2023).

Because decision-making is understood to be a continuous process in which incoming information is constantly evaluated, updated, and integrated (DasGupta et al., 2014; Groschner et al., 2018), studies using stationary decision assays often fit decision outcomes and reaction times to investigate the determinants affecting decision-making processes (e.g., Gold & Shadlen, 2007; Ratcliff & McKoon, 2008; Shadlen & Kiani, 2013; Urai et al., 2019). The continuous nature of the decision process is even more apparent in decisions on the move, which require decision-makers to carry out their decisions while simultaneously processing information and adjusting their movement trajectory accordingly. As such, it may be possible that upcoming decisions on the move are predictable using trajectory and other kinds of movement information. Indeed, recent studies make clear that interactions between locomotor behavior and spatial features within trials can reflect signatures of upcoming decisions, and possibly read out internal decision dynamics (Pinto et al., 2018; Sridhar et al., 2021; Kane et al., 2023; Molano-Mazón et al., 2024). These findings challenge the notion, implicit in the terminology generally employed in Y-maze studies, that decisions happen at *points*, rather than extend through time and space.

Here, we use the Y-maze to study the dynamics of locomotor decision-making in *Drosophila melanogaster*. Operationally, we consider two kinds of decision-making: Locomotor decision-making, the continuous process of choosing among possible movement behavior expressions (see Molano-Mazón et al., 2024), and turn-decisions, discrete spatial outcomes arising from integrated locomotor decisions (see e.g., Bett et al., 2012; Thomas et al., 2014). Specifically, we examine flies making continuous locomotor decisions while walking that add up to produce discrete turn-decisions at the maze intersection. We show that upcoming turn-decisions can be often predicted from locomotor behavior. The evolution in predictability exhibits distinct phases across both spatial and temporal domains. While predictability increases gradually over time, we can map sharp increases in spatially-dependent predictability to distinct locations within the maze. We further provide evidence that these dynamics depend on sustained locomotor dynamics, which suggests an active process like working memory (Fontana et al., 2019; Cleal et al., 2020; Cleal et al., 2021; Redhead et al., 2023). Moreover, we show that the evolution of predictability in the spatial domain differs across sensory and information-processing mutants, indicating that spatiotemporal decision-making is dependent on intact visual, mechanosensory, and dopaminergic circuitry. Last, we demonstrate that human trajectories through virtual versions of our Y-maze experiment exhibit space-varying decision predictability that is strikingly similar to flies, suggesting our findings may generalize across animals.

## Results

### Hints of future turn-decisions

To study the spatiotemporal dynamics of turn decisions in flies, we employed a symmetrical Y-shaped maze (Fig. 1A). Individual flies were placed in separate mazes and allowed to walk freely for a duration of 2 hours. We tracked the trajectories of each fly throughout the experiment (Fig. 1A-B, example fly). During their time within the maze, flies made multiple turn-decisions. That is, upon reaching the intersection, flies faced the choice of turning either to the unoccupied arm on their left or the unoccupied arm on their right. Throughout the experiment, each fly made numerous such turn-decisions, which allows for the study of their location-dependent choice dynamics over time.

**Figure 1.**
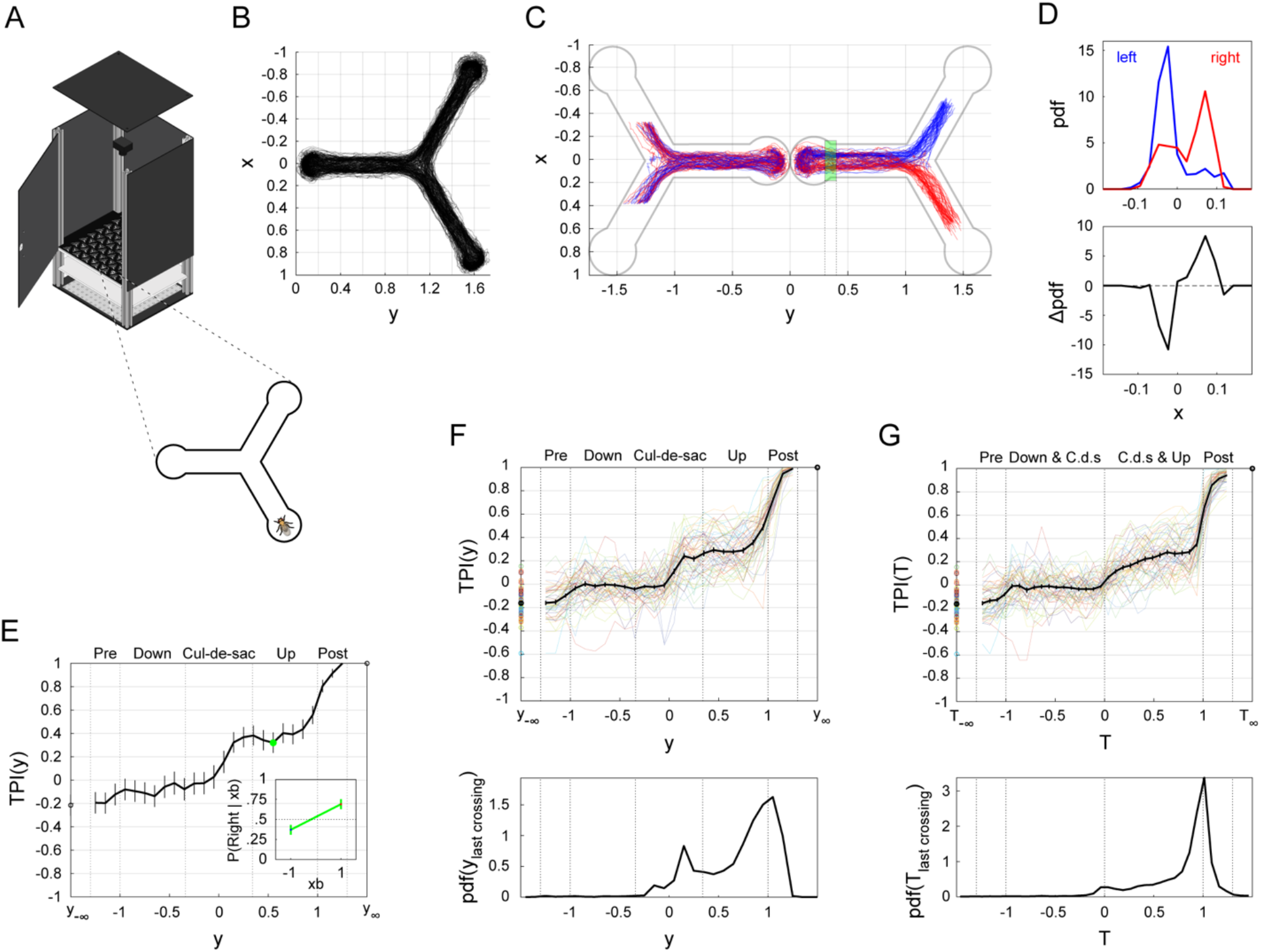
Measuring fly turn predictiveness in a Y-Maze. (A) Schematic of the Y-maze assay. Left: Flies were placed into an array containing many individual Y-mazes and the positions of the flies were recorded for 2 hours. Right: Detail of Y-mazes containing individual flies. (B) Trajectories of an example fly over the experimental duration (2 hours). (C) Normalized trajectories for the fly shown in B. All turn-decisions made from the bottom arm were standardized by assigning negative values to movements towards the bottom cul-de-sac of the bottom arm, positive values to movements towards the intersection (away from the bottom cul-de-sac), and zero to the bottommost point of the cul-de-sac. Blue and red: turn-decisions made from the bottom arm to the left and right arms, respectively. (D) Emerging indicators of turn predictability. Top: *PDF*(*x* |0.3 ≤ *y* ≤ 0.4, *Turn direction*), probability density functions of *x*-locations, conditioned on turn direction and *yRange* = {0.3 ≤ *y* ≤ 0.4} (green area in C). Color coded as in C. Bottom: *ΔPDF*(*x* |0.3 ≤ *y* ≤ 0.4); the difference between these two PDFs emphasizes that predictability of upcoming turn-decision could be read from x-locations within specific *y*-ranges. (E) Turn Predictiveness Index (TPI) in the spatial domain for the example fly in B-D. Black line denotes example TPI curve across all *yRange*s. Green circle denotes TPI value for the example *yRange. TRI*(*yRange*) measures how predictable a fly’s upcoming turn direction is with respect to its average x location. *TRI*(*yRange*) is given by the difference between two conditional probabilities: the probability of turning right, given that the average *x*-location is towards the right of the horizontal midline and the probability of turning right, given that the average *x*-location is towards the left (eq. 1 & Materials and Methods). Inset: Illustration of the TPI derivation for the example *yRange* (green circle, green area in C). In the example range, the fly turned right in 74% of trials in which its average x-location was to the right, but turned left in only 36% of trials in which its average x-location was to the left. Thus, the TPI value for the example *yRange* is 0.38. Error bars denote Standard Error (SE). (F) Average TPI curve in the spatial domain and corresponding PDF of last midline-crossings. Top: Average TPI curve (black) in the spatial domain over the entire sample (*n* =55), overlaid on the TPI curves of individual flies. Error bars denote Standard Error of the Mean (SEM). Bottom: PDF of last midline-crossings (LMCs), *PDF*(*y* |*LMC*), across all trials and flies (also see: Fig. S1C for the *PDF*(*y* |*LMC*) of individual flies). LMC is defined as the last *y*-location within a given trial in which the *x*-location changed polarity (i.e., last crossing of the horizontal midline). (G) As in F, for the temporal domain. The two tail values of the TPI curves in E-G correspond to the asymptotic behavior of the TPI before and after the fly entered the bottom arm, respectively. The value of 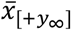 (resp. 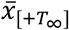) is determined by the eventual turn-decision, resulting in the TPI curve saturating at 1 (Eq. 1). With 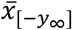 (resp.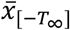) determined by the previous arm. TPI in the temporal domain for the example fly in B-E can be found in Figure S1A.

To gain insights into the movement patterns of flies as they navigate through the maze, we standardized the trajectory data of all turn-decisions made from the bottom arm (Fig. 1C). We considered a trial to be all of the movements a fly made from the point of entering the bottom arm, to turning around at the “cul-de-sac”, to walking up the bottom arm, and finally exiting to either the left or right arm. Prior to reaching the cul-de-sac, trajectories leading to right or left turn-decisions appear indistinguishable. However, trajectories that eventually resolve as right or left turns begin to diverge as the fly progresses away from the cul-de-sac and towards the intersection. Specifically, the fly appears to move more towards the right when subsequently turning to the right arm (*x* >0) and more towards the left when later turning to the left arm (*x* < 0). Importantly, the *x*-location of a fly does not deterministically predict the upcoming turn direction prior to the moment the turn is scored; i.e., a fly walking along the left side of the bottom arm wall may eventually make a turn into the right arm, and vice versa. Nevertheless, within a given *y*-range (i.e., a segment of the arm), *x*-locations could be predictive of the upcoming turn direction. For example, considering values of *y* between 0.3 and 0.4, *yRange* = {0.3 ≤ *y* ≤ 0.4}, the probability density functions (PDFs) of all *x*-locations markedly differ between trajectories of eventual left and right turns (Fig. 1D). These turn direction-dependent differences in flies’ *x*-locations allow for closer examination of trajectory-dependent turn decision predictability.

To measure turn-decision predictability in different spatial areas of the maze, we devised the Turn Predictiveness Index (TPI). This index measures how predictable a fly’s upcoming turn-decision is based on the average *x*-location, while encoding the concordance between that location and the left-right side of the eventual choice in its sign. We computed the TPI for each interval [*y*_1_, *y*_2_] along the y-axis of the maze as follows: 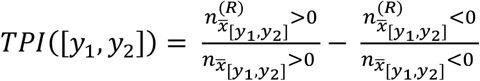 where 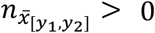 is the number of trials where the average *x*-position of the fly, while in the [*y*_1_,*y*_2_] interval, is greater than 0 (on the right), 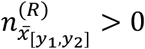 is the number of such trials that eventually resolve in a turn to the right, and the second ratio is similarly defined for trials where the average *x*-position is less than 0 (on the left). Thus, for each interval [*y*_1_, *y*_2_], TPI can range from −1 to 1, where its magnitude is a measure of predictability about the turn-direction. For example, for 0.3 ≤ *y* ≤ 0.4 (the *yRange* in Fig. 1C, green & 1D), the fly turned right in 69% of the trials in which its average *x*-location within this *yRange* was to the right, but in only 37% of the trials in which its average *x*-location within this *yRange* was to the left (Fig. 1E, inset). Hence, the TPI value for that example fly in that *yRange* is 0.69 −0.37 =0.32 (Fig. 1E, inset, green). Thus, the TPI provides a simple, signed point measure for predictability in each *y*-bin, and thus allows us to quantify how turn-decision predictability evolves as the fly progresses through the maze (Fig. 1E).

When calculated across the spatial extent of the maze, the average TPI across flies (Fig. 1F, top, black; n = 55) exhibits distinct phases, with sharp increases just before the maze arm intersection around *y* = 1 (not surprisingly: this is where turns are scored as left or right), but also in the cul-de-sac. Between these sharp increases are phases where the TPI remains relatively stable. The first value of this curve (*TRI*(*y*_−∞_)) captures the degree to which the current decision depends on the previous decision (sequential effect) and *TRI*(*y*_∞_), by definition, equals 1, since the x-position at that point defines the turn outcome. Taken together, the TPI metric captures core behavioral determinants of locomotor dynamics in movement decisions.

Locomotor decision studies often set a spatial threshold for recording the decision outcome in each trial (e.g., Buchanan et al., 2015; Dylda & Wang, 2022). While operational definitions as such are useful, their implementation effectively disregards the continuous nature of movement trajectories and their interaction with the emerging decision outcome. Indeed, distinct phases evident in the TPI curve (Fig. 1E) imply a dynamic decision-making process (also see Sridhar et al., 2021), in which locomotor decisions are made continuously as the fly traverses the maze and interacts with its geometry, with increased sensitivity to particular regions in space. This is reminiscent of ramping activity that is associated with static decision-making, in which neural activity prior to the actual decision becomes with time more and more predictive of the outcome motor action (Gold & Shadlen, 2007; Shadlen & Kiani, 2013; Gorbonos et al., 2024).

Additional evidence supporting this notion can be found in the midline-crossings carried out by the flies as they traverse the bottom arm. During the traversal of the maze in a given trial, flies make multiple midline-crossings, i.e., they move from one side of the horizontal center of the maze to the other (Figs. S5A & S1B). The distribution of the y-position at the last time the x-position crosses the maze midline, which determines the outcome of the future turn-decision (Fig. 1F, bottom) reveals multiple peaks, particularly in regions where the TPI sharply increases (Fig. 1F, top). This is apparent both when considering the entire sample (Fig. 1F, bottom), as well on the level of individual flies (Fig. S1C). If decisions were consistently made at a single region of the maze, we would instead have expected a concentrated, unimodal distribution of midline-crossings. In contrast, the multimodality of the midline-crossing distribution supports the conclusion that decisions are not confined to a single, critical location but are, instead, distributed over space in ways that are influenced by specific maze features.

### Future turn-decisions can be predicted well before a fly reaches the maze intersection

We asked if the TPI also evolves in phases over time, the way it evolves over space. We computed *TRI*(*T*), the TPI over relative time bins (Fig. 1G, top). Similar to TPI in the spatial domain, we define *T* = −1,0, 1 as the time points when the fly enters the bottom arm, reaches its minimal location, and leaves the bottom arm, respectively. Unlike TPI in the spatial domain, *TRI*(*T*) of each time bin is not strictly confined to a particular region of the maze. Instead, this binning method allows for the quantification of the dynamics of predictability in the temporal domain, separately for downward and upward traversals of the bottom arm. Similar to the spatial domain, the average *TRI*(*T*) curve displays distinct phases of increase in predictability: the initial above-zero increase occurs when flies switch from downward to upward motion (*y* =0), while the most abrupt increase occurs just before flies reach the intersection. Compared to the average *TRI*(*y*), the average *TRI*(*T*) increases more gradually (Fig. 1G). This occurs because flies slow down when switching from downward to upward motion within the cul-de-sac, spend relatively more time within this region, with longer path-lengths (Fig. S1D-E). As this kinematic sensitivity to specific maze regions is reduced over temporal binning, the rate of change in predictability during upward motion appears smoother in the temporal domain.

### Future turn-decision symmetry often breaks within the cul-de-sac

We were intrigued by the sharp increase in TPI in the cul-de-sac (Fig. 1F-G) and examined fly behavior in this region in more detail by computing the average velocity vector field within the cul-de-sac (Fig. 2A). As flies approach the edge of the cul-de-sac (*y* < 0), their average motion is largely symmetrical from left-to-right. However, as they progress away from the edge (*y* >0), this symmetry is broken and lateralized patterns of motion emerge for trajectories culminating in left turns (Fig. 2A, top) versus right turns (Fig. 2A, center). In both of these cases, the average motion out of the cul-de-sac is biased toward the side of the arm of the eventual turn-decision. Thus, well before the actual turn takes place, distinctive motion patterns emerge within the cul-de-sac that predict the direction of the flies’ future turns (Fig. 2A, bottom).

**Figure 2.**
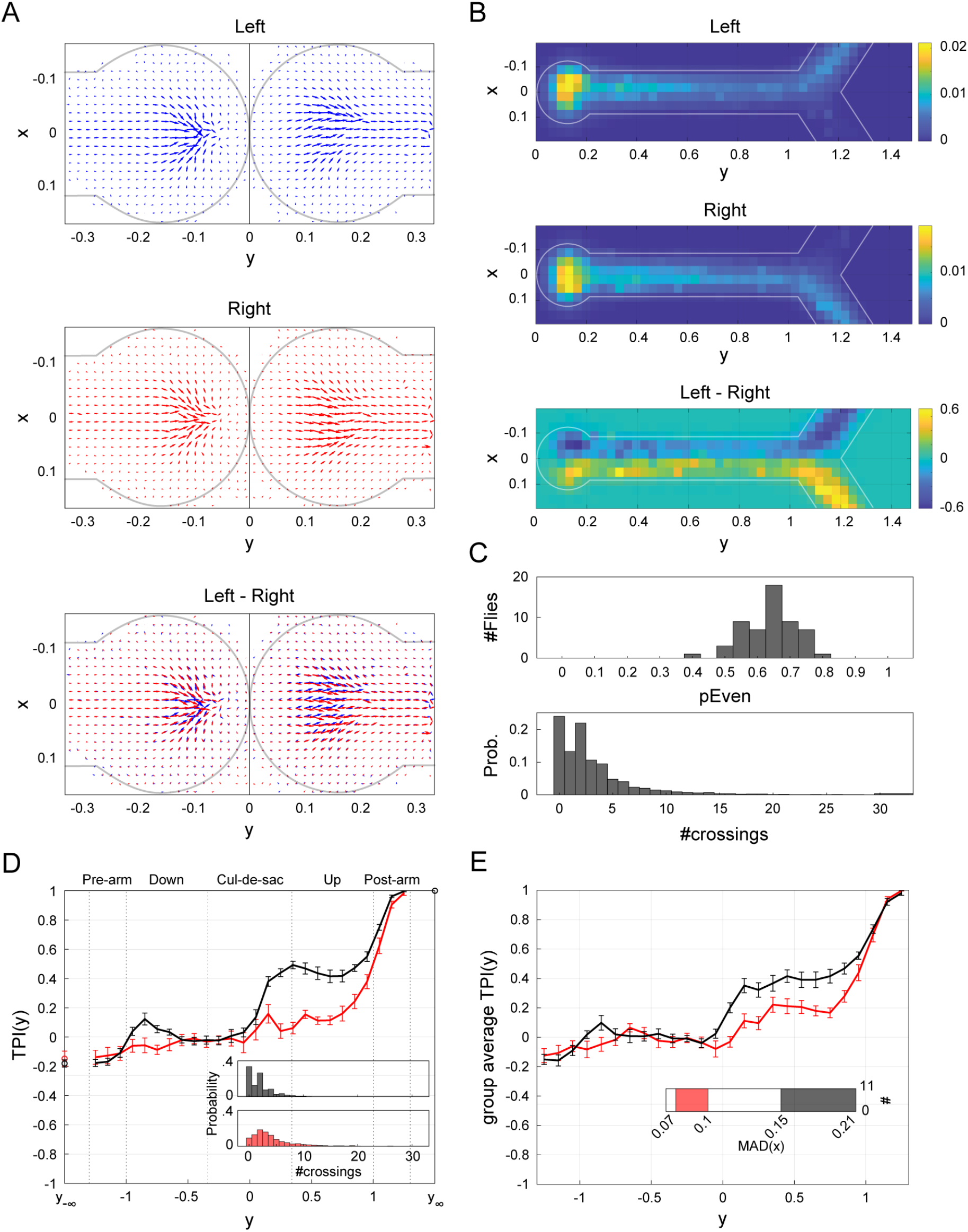
Locomotor dynamics and lateral tendencies. (A) Quiver plots depicting average motion within the cul-de-sac across flies (*n* =55). Quivers are computed separately for inward (*y* < 0) and outward (*y* >0) motion and separately for left (blue) and right (red) turns, based on flies’ corresponding velocity vectors (Supporting Information). In each 2D bin, the vector direction represents average motion direction across all flies, while the vector length indicates the relative frequency of visits to that bin (Supporting Information). Bottom: left and right turns, overlaid. Gray outline indicates approximate arena wall location. (B) Heat maps depicting the bivariate histograms for upward traversal (*y* >0), averaged over all files (*n* =55). Top: left turns-decisions. Center: right turn decisions. Bottom: right-left. Color temperature denotes probability (colorbar). White: representation of Y-maze outline. (C) Local lateral tendencies. Top: The distribution of *P*_*Even*_ over the sample (*n* =55). For each fly, *P*_*Even*_ denotes the fraction of trials in which the fly made an even number of horizontal midline-crossings (passes through *x* =0), #*crossings*, during upward traversal away from the cul-de-sac (*y* >0.34) until the end of the trajectory. Namely, 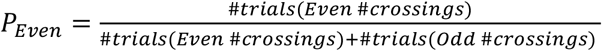. Bottom: The probability mass function of #*crossings*, for all trials (7904 trials). (D) Average TPIs for flies with lowest (red) and highest (black) *P*_*Even*_ values (calculated as in C, top; *n* = 11 per group). Inset: Probability mass functions of #*crossings* in each group (color coded as in the main panel; def. as in C, bottom). (E) As in D, for global lateral tendencies: Average TPIs for flies with lowest (red) and highest (black) MAD scores (*n* = 11 in each group). Inset: distribution of MAD scores over the sample. The score of each fly, *MAD*(*x*|0.34 < *y* < 1), computes the median absolute deviation from the horizontal midline for upward motion from exiting the cul-de-sac until the edge of the bottom arm. While these MAD scores correlate with the MAD scores corresponding to movement down the arm towards the cul-de-sac (*MAD*(*x*| − 1 < *y* < −0.34); Pearson *ρ* =0.53, *p* < .001), flies’ TPIs do not differ based on the latter (*MAD*(*x*| − 1 < *y* < −0.34) c.f., Figs. 2E & S2).

### Flies are non-deterministic wall-followers

The emergence of lateralized motion patterns within the cul-de-sac seeds a broken symmetry that propagates along the entire length of the arm, as is evident in the average occupancy of maze positions across flies and trials (Figs. 2B & 1E-G). Naively, this evokes a well-known bias of flies to follow walls (Soibam et al., 2012). However, this tendency is not deterministic, with TPI values less than 1 (∼0.2 to 0.4) for most of the arm beyond the cul-de-sac (see Fig. 1E-F, *y* >0.34 *&* Fig. 1G, *T* < 1) indicating that flies’ specific trajectories often diverge from the average patterns. Indeed, as they traverse the maze, flies often leave the wall they are following to cross the midline of the arm multiple times (3.7±2.1; mean midline-crossings after the cul-de-sac ± std) before making their final choice, even within the maze arm intersection (Figs. 1F, bottom & S1C). The parity of number of midline-crossings fully determines how a fly’s x-position relates to its future turn-decision: if the fly crosses the midline an even number of times, including 0, it will turn in the direction of the side it is currently on, and conversely if it crosses the midline an odd number of times, it will make a turn-decision in the opposite direction of its current position. Thus, the parity of midline-crossings provides a trial-specific measure of relationship between a fly’s position when it exits the cul-de-sac and its eventual turn-decision.

### Broken turn-decision symmetry often exhibits a working memory-like persistence

We define *P*_*Even*_ as the probability that #*crossings*, the number of midline-crossings through *x* =0 made after leaving the cul-de-sac in a trial, is even 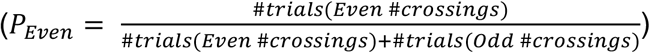. Across flies, *P*_*Even*_ is significantly biased toward making even numbers of midline-crossings per trial (two-sided Sign Test for *P*_*Even*_ −0.5, *p* < 0.001); 65% of flies exhibit *P*_*Even*_ significantly greater than 0.5 (Binomial test, *p* ≤ 0.023, two sided, not corrected for multiple comparisons). These results are in line with the observation that TPI values within the cul-de-sac are positive. That is, if an individual fly were equally likely to make an even number of even or odd midline-crossings after the cul-de-sac, its average *x*-location within the cul-de-sac would no longer predict the eventual turn-decision.

An implication of this analysis is that the TPI curves of flies with larger *P*_*Even*_ will display a sharper increase within the cul-de-sac compared to flies with smaller *P*_*Even*_. This is indeed what we find when comparing the average TPI of flies in the two extreme *P*_*Even*_ quintiles (Fig. 2D). Another prediction is that positive TPI reflects an excess of trials with even midline-crossings compared to odd midline-crossings. We examined the distribution of #*crossings* across all files after they leave the cul-de-sac (Fig 2C, bottom). As predicted, the frequency of 1 midline-crossing appears disproportionately lower compared to the frequency of both 2 and 0 midline-crossings.

The disproportionate occurrence of even amounts of midline-crossings after the cul-de-sac is at least partly due to excessive occurrence of zero midline-crossings, i.e., trials where flies follow a single wall all the way up the arm. However, the abundance of zero midline-crossings alone does not explain the observed bias of *P*_*Even*_ >0.5 in the sample; trials with one or more midline-crossing still have an even parity bias (Sign Test, two-sided, *p* =0.027). This significant tendency to make even #*crossings*, even when the fly does move away from the wall, suggests a form of memory of the lateral symmetry breaking that happened within the cul-de-sac, even through subsequent motion along the maze arm.

### Flies’ tendency to walk close to walls correlates with the predictability of their turn-decisions

Whereas *P*_*Even*_ reflects the effective commitment to a side after leaving the cul-de-sac on a trial-by-trial basis, we were also interested in quantifying the tendency to be close to walls irrespective of trial context. Specifically, we were interested in the general relationship between wall proximity and turn-decision predictability. We hypothesized that flies that remain closer to the walls of the arm will exhibit sharper increases in their TPI curves. To test this, we computed *MAD*(*x*|0.34 < *y* < 1), the median absolute deviation (MAD) from the horizontal midline of the x-position of the fly after it exits the cul-de-sac and before it reaches the intersection, for each fly across all trials. The average TPI curve of those flies with the largest *MAD*(*x*|0.34 < *y* < 1) values displays a substantially sharper increase within the cul-de-sac compared to the group with lowest values, in which TPI remains low until the intersection (Fig. 2E). Notably, while these MAD values do correlate with corresponding MAD scores computed on the movement *down* the arm (*MAD*(*x*| − 1 < *y* < −0.34); Pearson *ρ* =0.53, *p* < .001), flies’ TPI curves do not substantially differ when split according to the latter measure (Fig S2). Thus, the magnitude of turn predictability correlates specifically with flies’ tendencies to stay close to the wall *after* they exit the cul-de-sac.

### Wall-following alone cannot explain working memory-like patterns of locomotion

To isolate the contributions to turn-decision predictability of wall-following versus the working memory-like patterns seen in the parity of midline-crossing, we employed an agent-based modeling (ABM) approach and examined the Y-maze motion of agents with different locomotor rules, including variation in their tendency to follow walls (Fig. S13). We found that agents whose movement was Brownian motion-like, or who only followed a random walk in their heading, exhibited TPI curves that failed to capture the long-range predictability of turns (Fig. S13A). Adding a wall-following tendency to their locomotion rules extended the predictability of turns as far back as the cul-de-sac. However, these agents only showed increases in TPI within the cul-de-sac at implausibly high levels of wall-attraction (Fig. S13A, F), and never exhibited a bias towards even-parity beyond zero midline-crossings (Figs. S13E, left & S15C). This modeling supports the conclusion that while wall-proximity correlates with turn predictability, wall-following (as a locomotor mechanism) cannot recapitulate the spatial patterns of turn predictability and working-memory like local trial structure seen in real flies.

### Patterns of turn-decision predictability are invariant to maze arm length

Taken together, the above results suggest that flies do not necessarily behave as a “simple” particle guided by simple rules. Instead, they may exhibit more complex capacities, actively using information about their previous positions to guide their movement away from the cul-de-sac. However, it is still possible that these results are engendered by a passive process, wherein a “memory” of the motion within cul-de-sac arises from a spatially-imposed interaction between static statistics of fly movement and the particular length of the arm. Under this passive hypothesis, the short maze arms do not give flies enough time or space to render their ultimate turn-decision independent of their position when exiting the cul-de-sac. Said another way, the short arm length may not allow a propagating (perturbation-like) “memory” of the movement within the cul-de-sac to be averaged out (forgotten) during movement away from the cul-de-sac. Under this hypothesis, predictability within the cul-de-sac will decrease as the distance and time to the intersection increases. That is, because the memory of position within the cul-de-sac is given more space and time to average out in a longer maze. Alternatively, if predictability in a longer maze remains comparable to that of the short maze, then flies would appear to actively retain information about their preceding motion within the cul-de-sac. We tested these predictions by examining the locomotor behavior of flies in a Y-maze with arms twice as long as those in the original, short maze (Fig. 3A).

**Figure 3.**
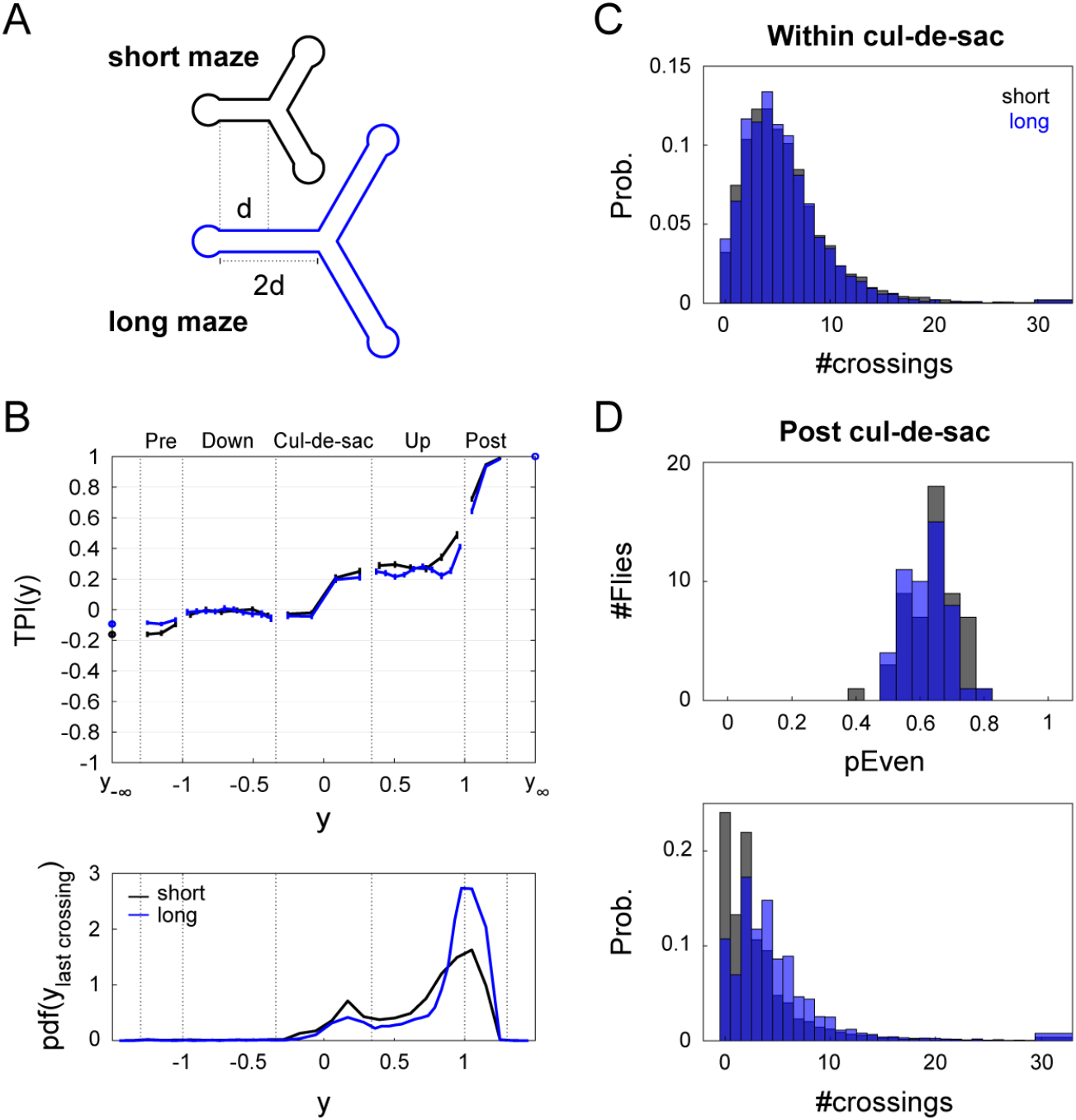
Cul-de-sac motion signature is maintained in a longer maze. (A) Details of Y-mazes with short (black) and long (blue) arms. The longer Y-maze has arms twice as long as those in the short maze. (B) Average TPI curve in the spatial domain and corresponding PDF of last midline-crossings. Top: Average TPI curve in the spatial domain of flies in the short (black, *n* =55) and long mazes (blue, *n* = 50). Error bars denote Standard Error of the Mean (SEM). Bottom: PDF of last midline-crossing (LMC), *PDF*(*y* |*LMC*), across all trials and flies. Bins ensure spatial alignment of maze regions (cul-de-sac, up or down the arm, post arm; see Fig. S3B-C for absolute and relative bins). (C) The probability mass function of #*crossings*, the number of horizontal midline-crossings **within** the cul-de-sac, for all bottom trials in the short (7904 trials) and long Y-mazes (7360 trials). (D) Local lateral tendencies post cul-de-sac. Top: The distribution of *P*_*Even*_ over the two samples.*P*_*Even*_ denotes the fraction of trials in which the fly made even #*crossings* during its upward traversal **away** from the cul-de-sac (*y* >0.34) until the end of the trajectory. Bottom: The probability mass function of #*crossings* **post** cul-de-sac, for all bottom trials in the short and long Y-mazes.

As expected, the time flies spend walking up the arm in the longer maze is significantly longer than in the short maze (median: 2.7 and 1.7 seconds, respectively; Wilcoxon rank sum test, *p* < 0.001; Fig. S3A, right). When we compared the average TPI curves of flies in the short and long mazes (Fig. 3B Top), we found that, contrary to the expectation under the passive process hypothesis, TPI curves in the short and long mazes are remarkably similar, both within the cul-de-sac (−0.34 < *y* < 0.34: Figs. 3B & S3B) and outside of the cul-de-sac (Figs. 3B & S3B-C), as well as in the temporal domain (Fig. S4). We observed that within the cul-de-sac, flies made similar numbers of midline-crossings in both mazes (mean±std: 5.8±1.5, 6.2±2.3, long and short, respectively; Fig. 3C). Conversely, flies in the long maze made 40% more midline-crossings after leaving the cul-de-sac (mean±std: 5.2±1.9, 3.7±2.1, long and short, respectively; Fig. 3B, bottom). Consistent with the similar TPI curves, we found a similar, significant enrichment (Sign Test, two-sided, *p* < .001) for flies making an even number of midline-crossings in the long maze compared to the short maze (Fig 3D, top). Most flies (94%) exhibited *P*_*Even*_ >0.5 and 70% of the flies had a *P*_*Even*_ value that was significantly larger than 0.5 (Binomial test *p* ≤ 0.0482, two sided, not corrected for multiple comparisons). Thus, there is no evidence that increasing the arm length decreased the predictability of future turn-decisions by allowing the flies more space and time to fluctuate away from the side of their symmetry that was broken in the cul-de-sac. Notably, this occurs despite the fact that flies make more midline-crossings in the long maze arms. While flies in long mazes make proportionally fewer trials with 0 or 2 midline-crossings, they make proportionally more with 4, 6, 8 etc. (Figs. 3D, S3B-C, & 2C-D), rendering the net turn-predictability comparable between maze geometries. Finally, we also considered the possibility of a third hypothesis, under which flies’ positive TPI enriched *P*_*Even*_ is due to a coincidental alignment of a characteristic locomotor curvature wavelength and length of the maze arm (as well as any maze arm that is an integer multiple longer). There are multiple reasons to believe this is not the case. First, distributions of #*crossings* of individual flies in the long maze exhibit multiple peaks at even numbers, suggesting parity tendency rather than a tendency to move in particular wavelengths (Fig. S5E). Second, if we count the number of midline-crossings that occur between the exit of the cul-de-sac and continuously-varying positions between the lengths of the short and long arms, parity tendencies are preserved (Fig. S5D). This is also true considering synthetic mazes shorter than the short maze. Third, when we synthetically produce crossing sequences by sampling from all inner-crossing-intervals of each fly in the long maze, thereby disrupting any dependence between consecutive crossing intervals, we fail to find any parity tendencies in either the long or short maze (Fig. S5B-C). Thus, even midline-crossing bias does not seem to emerge naturally by the interaction of a fly turning wavelength and maze arm length. Thus, taken together, the observations above are consistent with a working memory-like mechanism that persists long enough to extend the predictability of future turn-decisions twice as far in space.

### Disrupting sensory and circuit genes increases the predictability of turn-decisions

To explore mechanisms underlying turn behavior predictability, we examined the behavior of flies with mutations affecting their sensory, and circuit functions. Specifically, we measured behavior of flies mutant in genes encoding the mechanosensory channel NOMPC (Yan et al., 2012), the phospholipase-C NorpA (Bloomquist et al., 1988), dopamine Receptor 1 (Kim et al., 2003; *dumb*), and the transcription factor FoxP (Weigel et al., 1989). Respectively, these mutations are reported to affect gentle touch (Yan et al., 2012) and locomotor activity (Cheng et al., 2010), eliminate all vision (Pak et al., 1970), impair plasticity in the mushroom body (Kim et al., 2007) and the premotor central complex (Kottler et al., 2019), and affect locomotion and landmark fixation (Palazzo et al., 2020) and decision-making (DasGupta et al., 2014).

We found that mutant flies’ TPI curves, like wild type (WT) TPI curves, start increasing sharply within the cul-de-sac, remaining relatively stable as flies traverse up the arm, and exhibiting another substantial rise close to the intersection (Figs. 4A & S8). However, TPI(y) of the *dumb*^*2*^, *norpA* and *nompC* mutants significantly deviate from that of WT, even within the cul-de-sac (Fig. S6A, left). The *foxP* TPI curve did not significantly differ from WT. The higher predictability of the other mutants at the end of the cul-de-sac could neither be accounted for by higher predictability magnitude when entering the cul-de-sac, nor by sequential effects (Fig. S6A, center and right, respectively). Even though the *foxP* TPI curve did not differ from WT, we did observe that these flies were slower than WT, spending more time in all maze regions (Fig. S6B), and yet this change in dynamics did not translate into a change in turn-decision predictability.

**Figure 4.**
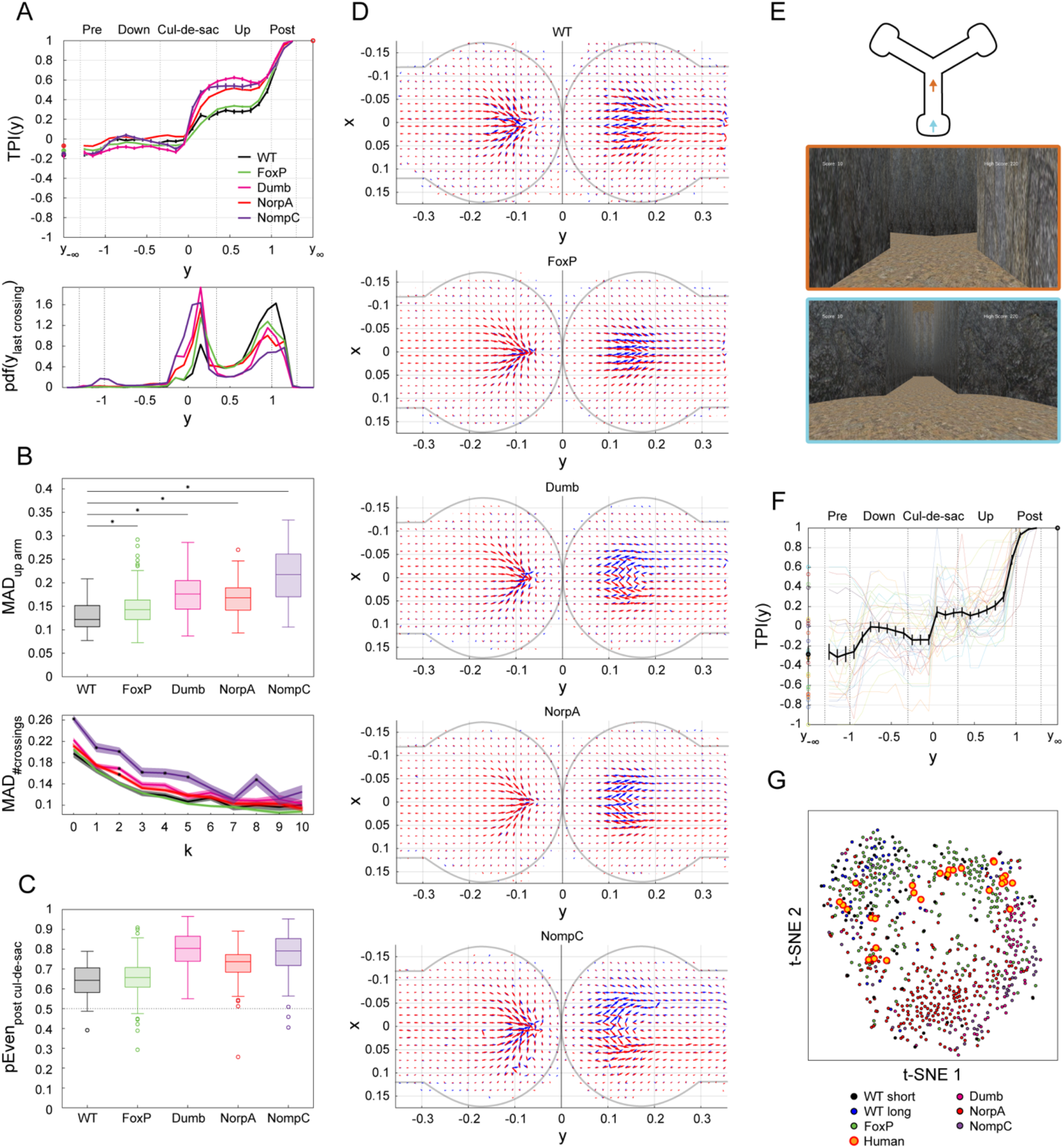
TPI in mutants and humans. (A) Average TPI curves in the spatial domain and corresponding PDFs of last midline-crossings across genetic lines: WT (black, *n* =55), FoxP (green, *n* = 243), dumb (magenta, *n* = 90), NorpA (red, *n* = 247) and NompC (purple, *n* = 54). Top: Average TPI curve in the spatial domain of flies in each line. Error bars denote SEM. Bottom: PDF of last midline-crossing (LMC), *PDF*(*y* |*LMC*), across all trials and flies in each line. (B) Global lateral tendencies. Top: Box plot depicting *MAD*(*x*|0.34 < *y* < 1) scores (computed as in Fig. 2E, inset) for each sample. For each box, the central mark indicates the median, bottom and top edges of the box indicate the 25th and 75th percentiles, whiskers extend to the most extreme data points not considered outliers, and outliers are plotted individually using symbols. Black horizontal lines: two-sided Wilcoxon Rank sum test for the difference in medians in MAD scores of mutant vs WT flies: *p* < 0.001, corrected for multiple comparisons, asterisks denote significance. Bottom: MAD values conditioned on the number of midline-crossings post cul-de-sac, *MAD*(*x*|0.34 < *y* < 1, #*crossings* = *k*), for each line. Lines: mean. Shaded area: SEM. Asterisk: significant difference between WT and mutant (two-sided Wilcoxon Rank sum test for the difference in medians of *MAD*(*x*|0.34 < *y* < 1, #*crossings* = *k*) scores of mutant vs WT flies, corrected for multiple comparisons). (C) Local lateral tendencies. Box plot (as in B, bottom) depicting *P*_*Even*_ across each sample. (D) Quiver plots depicting average motion within the cul-de-sac across flies in each line (as in Fig. 2A, bottom). (E) Schematic of the computerized version of the Y-maze task. Humans (*n* = 30) navigated a cave-like structure to collect hidden coins. Top: Schematic of the computerized Y-maze. Bottom: Screenshots of participants’ view at 2 example locations in the maze (frame colors correspond to arrows in Top). (F) Average TPI curve in the spatial domain for human participants (black), overlaid on the TPIs of individuals. Error bars: SEM. (G) Dimensionality reduction of the individual TPI curves in the spatial domain of humans and all fly lines via t-distributed Stochastic Neighbor Embedding (tSNE). Each point represents the TPI of an individual decision-maker between −1 ≤ *y* ≤ 1 and distances between the points approximate the cosine similarity among the TPIs.

We previously saw that flies with on average greater wall proximity had greater turn-decision predictability (Fig. 2E). This pattern held with the *dumb, norpA* and *nompC* mutants (Fig. S11C), which exhibited significantly greater wall proximity compared to WT (as measured by the MAD of their x-positions; Fig. 4B, top; two-sided Wilcoxon Rank sum test: p < 0.001). Considering the parity of midline-crossings after the cul-de-sac, these 3 mutant lines also make more even midline-crossings than WT flies (Fig. 4C; % even #*crossings*±SE: 65.1±0.54%, 80.5±0.36%, 73.4±0.19%, 79.1±0.45% for WT, *dumb, norpA* and *nompC*, respectively). Thus, the relationships seen between wall proximity and even parity midline-crossing and turn-decision predictability among different flies within the WT genotype also hold between WT and *dumb, norpA* and *nompC* mutant flies.

Examining the movement statistics of these lines in detail, we found that these mutants exhibit comparable numbers of midline-crossings within the cul-de-sac compared to WT flies (Fig. S6C, left mean±std #*crossings*: 5.8±4.7, 5.4±5.0, 4.9±4.1, 4.3±4.1 for WT, *dumb, norpA* and *nompC*, respectively). In contrast, the three mutant lines made fewer midline-crossings after the cul-de-sac (c.f., Figs. 4A, bottom & S6C, right). In fact, the larger *P*_*Even*_ values observed in these mutant flies (Fig. 4C, Fig. S6C, right) primarily stem from the excessive occurrence of zero midline-crossings after the cul-de-sac (Fig. S6D; %trials with no midline-crossings ±SE after the cul-de-sac: 24.0±0.48%, 50.9±0.45%, 40.4±0,21% and 57.2%±0.54% for the WT, *dumb, norpA* and *nompC*, respectively). When excluding trials with zero crossings, TPI curves of mutants appear much more similar to that of WT flies; multiple measures of turn-decision predictability have no statistically significant difference between WT and nompC flies, for instance, if zero-crossing trials are excluded (Fig. S9). Therefore, the main effect of these mutants in increasing turn-decision predictability seems to stem from an increased rate of trials in which mutant flies walk from the cul-de-sac to the intersection, following one wall the entire way.

The behavior of the mutants with higher turn-decision predictability within the cul-de-sac reveal motion patterns that are more spatially separable in trials ending with left versus right turn directions (Figs. 4D, S7, & S10). Taken together, the choice dynamics exhibited by WT and mutant flies hint at nuanced decision-making processes that vary across genetic lines and highlight the importance of different sensory modalities flies use when making decisions. Specifically, vision (NorpA), mechanosensation (NompC), as well as working memory (Dop1R1) appear to mediate the interaction of flies’ probabilistic locomotor rules and their spatiotemporal environment.

### Future turn-decisions are predictable in other experimental contexts and humans

To assess whether future turn-decisions are predictable over space and time in other experimental contexts, we examined data from a previous study involving WT flies navigating a circular arena towards identical, equidistant targets (Sridhar et al., 2021). Using a measure equivalent to x-position as the predictor of future turn-decisions, we computed TPI curves from this dataset (Fig. S12). Despite the different arena geometry and task context compared to the Y-maze, flies displayed distinct phases of predictability increase in advance of when turn-decisions were officially scored, suggesting that choice dynamics extending over space and time is not likely to be a result of maze geometry alone. Specifically, TPI analyses appear to be applicable across diverse motion decision tasks.

Having seen that flies of different genotypes and in different choice geometries all exhibit turn-predictability that extends over space and time (while differing in the locomotor basis of this predictability), we wondered whether this might be a phenomenon that translates to humans. Fabricating Y-maze assays at a proportionate scale for humans and matching the throughput of our fly experiments is not feasible, so we recorded the virtual behavior of human participants engaged in a computerized version of a Y-maze task (Fig. 4E). Calculating TPI curves from their trajectories, we observed distinct phases akin to those found in flies (Fig. 4F). Unlike flies, human TPI curves exhibit a stronger sequential effect (also recognizable in more extreme TPI values at −*y*_∞_). Still, the overall TPI curve phases were similar for both flies and humans, especially for motion post cul-de-sac (c.f., Figs. 1F, top & 4F). Moreover, visualizing the TPI of all individual flies and humans in two dimensions using t-distributed stochastic neighbor embedding (tSNE) reveals that human TPI curves fall in similar clusters as wild type fly TPI curves. Thus, there may be deep homology in the predictability of turn-decisions across animals.

## Discussion

We have shown that locomotor decisions in a forced-choice arena environment are not made at a specific moment in time or at a specific point in space, but instead emerge dynamically over both space and time (Fig. 1). Using flies’ past locomotor features, we can predict their future turn-decisions much earlier than they occur and far from the region of the maze generally described as the “choice point.” Specifically, flies’ lateral location is a simple, strong predictor of eventual turn-decision. This predictability has characteristic spatial and temporal dynamics. Within the cul-de-sac located at the end of a maze arm, flies’ TPI exhibits a sharp sudden increase, which is followed by a more gradual rise, and culminates in a final increase as flies traverse the arm intersection.

Flies have a propensity to avoid open spaces and follow walls (Soibam et al., 2012). Thus, it is conceivable that the future turn-decision predictability patterns we observed were the result of simple locomotor rules interacting with the arena geometry. If decisions appeared ‘locked in’ simply because there was insufficient time and space to introduce independence in the locomotor pattern, we would expect lower turning predictability if this were remedied. This was not the case: When we lengthened the maze arm by a factor of two, we found that turn-decision predictability dynamics still have the same structure (Fig. 3). Specifically, while flies crossed the midline of the maze arms more often in the long maze arms (as expected), they also crossed the midline an even number of times proportionally more often. This pattern of locomotion meant that their position leaving cul-de-sac was just as predictive of their turn-decision, even though the intersection was farther away.

Taken together, these observations suggest that flies sustain a signature of their locomotion for relatively long periods of time, i.e., exhibit dynamics consistent with working memory. This is consistent with previous studies that have shown flies can guide their locomotion with spatial memories (Neuser et al., 2008; Ofstad et al., 2011; Kim & Dickinson, 2017). To investigate the possibility that memoryless locomotor mechanisms, interacting with the arena geometry might account for the predictability of turn-decisions, we implemented agent-based models of fly locomotion with tunable behavior rules (Fig. S13). Systematically increasing the degree of agent wall-following could account for high turn-decision predictability, but not the patterns of even and odd midline-crossings found in real flies. We also confirmed through resampling analyses that the empirical parity bias of midline-crossings was present throughout the maze arms and disappeared when the bout-by-bout structure of mid-line crossings was scrambled. We thus conclude that spatiotemporal dynamics of turn-decision predictability are not simply a byproduct of memoryless wall-following tendencies, but persist even when flies are given more time and space to uncouple their behavior in the cul-de-sac from their eventual choice outcome at the intersection.

We found that spatial predictability dynamics differ between WT flies and mutants with disrupted vision (*norpA*), mechanosensation (*nompC*), and dopaminergic signaling (*dumb*) (Fig. 4). Compared to WT, turn-decisions of *norpA* and *nompC* flies are significantly more predictable already within the cul-de-sac. This is driven primarily by these mutants frequently traversing from the cul-de-sac to the intersection without making a single midline-crossing (Fig. S6). It is plausible that flies rely on visual Information about maze arm locations as well as mechanosensory information about their distance from the wall to guide their locomotion (Cheong et al., 2020). In the absence of such sensory information, turn-decisions may be under fewer modifying influences (Pak et al., 1970; Yan et al., 2012) and thus more predictable.

We further show that turn-decision predictability of flies with disruptions to the Dop1R1 showed increased predictability like *norpA* and *nompC* mutants. Turn-decisions of *dumb* mutants are more predictable than those of wild type flies, even when trials with no-midline crossings are excluded (Fig. S9). Impairments in dopamine signaling pathways, especially in the mushroom body and central complex neuropils, have previously been implicated, across multiple species in altered salience-based decision-making (Zhang et al., 2007), locomotor action-selection (Kottler et al., 2019; Fisher et al., 2022), and spatial working memory (Mohandasan et al., 2020). *FoxP* mutants that have previously been reported to exhibit deficits in locomotor turning behavior, olfactory decision-making dynamics, learning deficits, and reduced activity (Groschner et al., 2018; DasGupta et al., 2014; Mendoza et al., 2014; Ehweiner et al., 2024) exhibited no substantial difference from WT flies in our assay. It is possible that our locomotor decision-making specifically depends on underlying neuronal processes that are independent of *FoxP*, or subject to compensatory regulation during development.

The higher predictability in these mutants reflects that the left-right symmetry they break in the cul-de-sac more often propagates all the way to the maze intersection. How symmetry breaks in the cul-de-sac is an open question. We can observe distinct locomotor patterns within the cul-de-sac that correlate with the eventual turn-decision (Fig. 2). However, it is not clear whether this reflects a process of decision-making itself, or a behavioral readout of a decision made prior to entering the cul-de-sac. It is possible, for example, that locomotor behavior within the cul-de-sac directly reflects accumulation of left-versus-right sensory evidence (Ratcliff & McKoon, 2008; Milosavljevic et al., 2010; Buerkle et al., 2021).

To assess whether spatiotemporal patterns of turn-decision predictability generalize across species, we asked human participants to virtually explore an online, 3D version of the Y-mazes (Cleal et al., 2020; Cleal et al., 2021; Redhead et al., 2023; Oliver et al., 2023). Human virtual locomotor behavior featured turn-decision predictability patterns that were, in many cases, qualitatively indistinguishable from flies (Fig. 4E-F). Specifically, despite the drastically different experimental setting and subjects, turn-decision predictability sharply increased when participants exited the cul-de-sac, and gradually increased until participants eventually turned through the intersection into another maze arm. The TPI curves of human Y-maze behavior cluster along with wild type fly TPI curves (from both short-and long-arm mazes), and are distinct from *norpA, nompC* and *dumb* mutant fly TPI curves in a t-SNE embedding (Fig. 4G). In summary, the predictability of Y-maze turn-decisions extends across space in similar ways in multiple species in different sensory contexts.

While TPI curves provide insight into turn-decision dynamics of flies and humans, the particular geometry of the maze is certainly contributing to the distinct phases of increasing predictability preceding choices. Indeed, TPI curves calculated for flies walking in open arenas exhibit different phases (Fig. S12). It is also worth emphasizing that while the locomotor trajectories of flies offer rich information about their decision-making processes, including future turn-decisions, interpreting these trajectories as direct readouts of internal decision processes is likely an oversimplification. Instead, they serve as proxies, seen through many distorting lenses such as biomechanical constraint and interaction with the environment, that hint at underlying decision dynamics. Studies tend to interpret locomotor decisions as happening at specific, critical locations and singular moments in time. Our findings challenge this notion by revealing, through TPI analysis, distinct phases of future choice predictability that extend over space and time. We conclude that turn-decisions in Y-mazes, and on-the-move decisions in other arena geometries, are not likely to be determined at a single moment in a single location. Rather, our results imply a dynamic decision-making process, where locomotor choices are influenced by multiple sensory systems and locomotor processes across different regions of the maze.

## Materials and Methods

### Fly lines and husbandry

Wild type flies used in behavioral assays were Canton-S. Mutant lines used were NorpA: norpA[EE5], NompC: nompC[b19]/SM6b, dumb2: w[1118]; Df(3R)Exel8159/TM6B, Tb[1], FoxP: Df(3R)by10, red[1] e[1]/TM3, Sb[1] Ser[1]. All fly lines were grown in bottles on Caltech formula medium at 25°C in temperature-controlled incubators on a 12h:12h light:dark cycle.

### Fly experiments

Flies were assayed at 23°C in acrylic Y-maze arenas organized in trays of 48 arenas. Trays were diffusely illuminated from the bottom. Prior to assay begin, flies were anesthetized using CO2 and aspirated into individual arenas. Flies were allowed 30 minutes recovery and exploration time before the experiment started. Over the experimental duration of 2 hours, flies freely moved through the Y-maze, and their positions and turns were recorded at 30 fps using a Blackfly GigE camera (PointGrey BFLY-U3-13S2M-CS) and tracked using custom MATLAB software (Werkhoven et al., 2019). Flies are tracked via their centroid location. Trays were illuminated using LED panels (Part BK3301, Knema LLC, Shreveport, LA). Due to considerable fisheye effect present in arenas at the tray periphery, trajectories of individual flies were manually inspected prior to inclusion and flies with tracking errors were excluded from the raw dataset (see Table S1, ‘post visual inspection’).

### Fly raw data and exclusion criteria

We retained trials that were made from the bottom arm to either the left or right arm. Because we were interested, for each turn from the bottom arm, in the trajectory of the fly from when it passes through the intersection to enter the bottom arm until it passes again through the intersection to exit the bottom arm and turn to either the left or right arm, we defined a bottom trial such that it includes these trajectories. To avoid using trials with missing trajectories, we excluded trials in which flies fail to reach the bottom part of the arm or trials that included missing values. Flies with fewer than 80 trials were excluded from the analyses (see Table S1, “post #trials threshold”). We also excluded outlier flies with trajectory bounding polygons that were misaligned compared to the sample and could not be corrected. Finally, we excluded individual trials that contained trajectory coordinates outside of the maze, but did not exclude additional flies based on this criterion.

### Human experiments

Ethical approval for this study was obtained from the University of Southampton Psychology ethics committee. Human participants (n = 50) completed an online, computerized version of the Y-maze (Fig. 4E; https://www.experiments.psychology.soton.ac.uk/mazes/ymcve_curved_r/). Participants were recruited via an online participant recruitment platform, prolific.co, in return for remuneration of £2. Participants were instructed to navigate a virtual Y-maze using the arrow keys on their keyboard and search for hidden coins in the cul-de-sac surroundings. The virtual Y-maze environment was scaled to simulate Y-maze arenas in which we assayed flies. Participants were instructed to collect as many coins as they could over the experimental duration (12 minutes).

### Human raw data and exclusion criteria

Initial exclusion of human participants from the dataset was based on minimal location across all trajectories. Participants were excluded if they did not reach the bottom edge of the bottom cul-de-sac. Overall, 15 human participants were excluded from analysis in this way. To maximize the number of analyzable trials, we rotated turns from the left or right (120 or 240 degrees) arm to resemble turns from the bottom arm. Of all resulting rotated trajectories, incomplete (e.g., not: left/right arm → bottom arm → left/right arm) or missing trajectories were excluded from analysis. Further, 5 participants with fewer than 28 valid trials were excluded from analysis.

### Analyses

#### Corrected trajectories

All analyses use turns made from the bottom arm to either the left or the right arm. That is, only valid bottom trials for the fly data and all valid rotated trials in the human dataset. These trajectories were then scaled and normalized. The scaling resulted in *y*′ = 1 denoting the upper edge of the bottom arm, just before the intersection, whereas *y*′ =0 denotes the minimal location reached during the experiment). X values were scaled proportionally to the y scaling, so that the maze x:y ratio did not change. To distinguish motion towards and away from the bottom arm’s cul-de-sac, the scaled trajectory of each trial *Tr* was then standardized according to the minimal *y*′ location in that trial,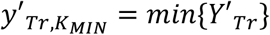. Specifically, *y*_*Tr,k*_ = −*y*′_*Tr,k*_ if *k* ≤ *K*_*MIN*_ and *y*_*Tr,k*_ =*y*′_*Tr,k*_ if *k* > *K*_*MIN*_. To reduce between-flies tracking shifts on the horizontal axis, the *x*-coordinates of each fly were centered (*x* → *x* −0.5 · (*max*{*x*| − 1 ≤ *y* ≤ 1} − *min*{*x*| − 1 ≤ *y* ≤ 1}) so that *x* =0 after centering reflects the horizontal midline of the bottom arm, over all of the fly’s valid trials. The centered *x*-coordinates were used in all analyses.

To capture behavior in the temporal domain, we also defined *t* as a variable denoting *relative* time spent in the bottom arm, separately for motion towards and away from the edge of the bottom cul-de-sac. Specifically, for each trial *Tr*, we used the value of *K*_*MIN*_ (see above) and identified the locations 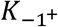 and 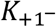, in which *y*_*Tr,k*_ >= −1 for the first time of and *y*_*Tr,k*_ ≤ 1 for the last time, respectively. We defined 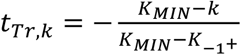 for *k* < *K*_*MIN*_,*t*_*Tr,k*_ =0 if *k* = *K*_*MIN*_, and 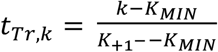 if *k* > *K*_*MIN*_. Thus, the resulting *t*_*Tr,k*_ values denote the relative time of downward (*y, t* < 0) and upward (*y, t* >0) motion within the bottom arm in trial *Tr*, where 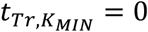 is time where the minimal *y* location in the trial is reached. A downward motion in the arm therefore spans from 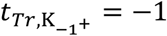 to 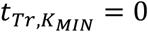 in the temporal domain (upward motion: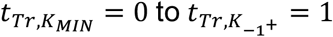).

#### Midline-crossings

The above centering allowed for straightforward identification of locations where midline-crossings occurred. Namely, a midline-crossing event in a trial is defined by *sign*{*x*(*k*) · *x*(*k* + 1)} = −1, and we define #*crossings*_*Tr*_ as the number of midline-crossings in trial *Tr* for a specific region in the maze - typically after exiting the cul-de-sac (e.g., Figs. 2C & 3D), but also within the cul-de-sac (e.g., Fig. 3C).

#### TPI - spatial and temporal

To assess the predictability about the upcoming turn-decision and its evolution in the spatial and temporal domains, we devised the Turn Predictiveness Index (TPI). This index measures, for each given spatial or temporal range, how predictive the horizontal (*x*) average location of a fly is about the evential turn-direction. In the spatial domain, the corrected trajectory data was parsed into 26 equal *y*-bins with ranges defined by *yRange*(*j*): = *Y*(*j*) ≤ *y* ≤ *Y*(*j* + 1) and *Y* = {−1.3,1.2, …, 1.3} (except for Fig. 3B, bottom, see caption). To derive the polarity of the horizontal location of the subject in trial *Tr* within the vertical location given by *yRange*, we defined *xb*(*yRange*)_*Tr*_: = *Singn*{ < *x*|*yRange, trial* = *Tr* >} (where < · > denotes average) as a binary variable whose value is −1 if the average *x*-location is toward the left and 1 if it is towards the right. The TPI of the subject in the bin given by *yRange, TRI*(*yRange*), is given by the difference between two conditional probabilities: the probability that the subject will turn right given that the horizontal location is *xb*(*yRange*)_*Tr*_= +1 and the probability that the subject will turn right given that the horizontal location is *xb*(*yRange*)_*Tr*_= −1 (eq. 2, equivalent to eq. 1). Thus, ranging from -1 to 1, the TPI magnitude (|*TRI*(*yRange*)|) quantifies how predictive *xb*(*yRange*) is about the upcoming turn-decision and the TPI polarity captures (in)congruency (i.e., whether or the subject turns to same *xb*(*yRange*) direction, *TRI*(*yRange*) >0 and *TRI*(*yRange*) < 0, respectively).

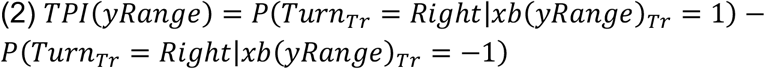

The two tail values of this curve, *TRI*(−*Y*_∞_) and *TRI*(+*Y*_∞_) reflect TPI (eqs. 1 & 2) at the first and last data points considered in each trial. That is, *xb*(−*Y*_∞_)_*Tr*_= −1 if the subject arrived at the bottom arm from the left arm and *xb*(−*Y*_∞_)_*Tr*_= +1 if they arrived from the right arm. Thus, with its value given by *P*(*Turn*_*Tr*_= *Right* | *Turn*_*Tr*−1_ = *Left*) − *P*(*Turn*_*Tr*_= *Right* | *Turn*_*Tr*−1_ = *Right*), *TRI*(−*Y*_∞_) measures the subject’s tendency to alternate between turn-directions, with positive (negative) values associated with repetition (alternation) tendencies. Similarly, *TRI*(+*Y*_∞_) captures the asymptotic behavior of the curve at the end of trajectory and it therefore saturates at 1.

The TPI in the temporal domain was computed similarly, only that we parsed the trajectory data into 17 equal relative time bins for *y* < 0 and 17 equal relative time bins for *y* >0, defined by *tRange*(*j*): = *T*(*j*) ≤ *t* ≤ *T*(*j* + 1), where *T*_*y* < 0_ = {−1.275,1.2, …, 0} and *T*_*y*>0_ = {0,0.075, …, 1.275}. We note that while the number of data points in a given spatial or temporal bin varies between trials, the number of data points in a given trial is roughly similar for temporal bins defined by *tRange*_-1 < y < 0_ and for temporal bins defined by *tRange*_0 < *y* < 1_, but could markedly differ for the spatial bins given by *yRange* (Fig. S1D-E, top).

## Data availability

All raw datasets and analysis scripts are available in the Zenodo archive at https://zenodo.org/records/13352340 (Lebovich et al., 2024). Data files and scripts are described in the read-me file.

## Code availability

The custom MATLAB codes used for all analyses are available in the Zenodo archive (Lebovich et al., 2024) and in the SpatiotempDM GitHub repository at: https://github.com/Lior-Lebovich/SpatiotempDM.

Python code for agent-based models can be found in the Zenodo archive and in the GitHub repository at: https://github.com/tomalisch/ABM-Drosophila.

## Acknowledgements

We thank Edward Soucy and Yuwei Li for their continuous help constructing assays. B.d.B. was supported by NIH/NINDS grant no. 1R01NS121874-01. Y.L. was supported by the Gatsby Charitable Foundation and by a grant from the DFG (CRC1080). Y.L. is the incumbent of the David and Inez Myers Chair in Neural Computation. I.D.C. acknowledge support from the Office of Naval Research Grant N0001419-1-2556, Germany’s Excellence Strategy-EXC 2117-422037984, the Max Planck Society, the European Union’s Horizon 2020 research and Innovation Programme under the Marie Skłodowska-Curie Grant agreement (#860949) and the PathFinder European Innovation Council Work Programme (#101098722).

## Supplementary Information

### Simulated agent-based fly models (ABM)

To investigate whether we could approximate fly decision-making in a Y-maze using only simple movement rules, we employed an agent-based modeling (ABM) approach. In this modeling framework, a simulated fly is treated as a circle of a constant radius, *r*_0_, traversing the maze and constrained by the surrounding walls. Its initial position is drawn from a distribution of valid positions, in which the entire circle lies within the maze. For each of the remaining steps, the position is updated based on a distribution of heading angles, speed, and distance to the surrounding walls of the maze. The simulated agent makes repeated turn-decisions, until the predetermined simulation duration is reached. We used scaled parameters that match the body size and speed of a WT fly in the short maze, as well as the framerate of the assay camera. We examined the behavior of different agents by implementing different position update rules on the distribution of heading angles and simulated multiple flies in each.

In its simplest form, which we use here as a baseline, a simulated agent in the above ABM framework follows a constrained Brownian motion. Specifically, the heading angle in each frame, which was matched to the sampling rate of the preceding behavioral assays, is drawn from a Uniform distribution (between 0 and 359) and the resulting direction is only constrained such that the circle is entirely confined by the maze walls. Angles are defined in such a way that 0 and 360 denote forward facing, and 180 denotes backwards. To further understand the motion choice dynamics exhibited by real flies, we extended the ABM by: (1) introducing heading angle choices depending on past angles correlated rather than random heading angles and, (2) incorporating a tunable parameter, *WF*, governing wall-following tendencies. In summary, for every frame f, simulated flies randomly pull a heading angle *α* from a Gaussian distribution of mean μ = 0, and variance ***σ***. The resulting heading angle is then used to update the last recorded heading direction. Next, simulated fly’s position is attempted to be updated based on the resulting heading direction and the fly’s speed s, which approximated real flies’ speed in mm/frame in the behavioral assays. If the proposed position would result in any part of the fly body circle to be inside the environment walls or outside of bounds, the closest valid position is instead chosen. Closest valid position is defined as the coordinate that (a) would result in the smallest current heading angle change and (b) would result in no part of the fly body circle to be inside a wall or outside of bounds. Wall attraction was set to occur to a degree given by *WF* whenever the distance, *d*(*t*), between the current position (centroid) of the simulated fly and the closest wall coordinate falls below a predefined fixed parameter *r*_*W*_ (*r*_*W*_ > *r*_0_; see Materials and Methods). Thus, for a given value of *WF* the agent updates its heading angle only based on (1), unless *d*(*t*) < *r*_*W*_, in which case the chosen heading angle is updated by the *WF*-weighted mean of the angle towards the closest wall and the current heading angle, as described in (2).

We simulated a first set of 3 experiments of increasing complexity using a combination of the simple rules described. In the first experiment, we simulated flies that followed Brownian motion. Every frame, heading angle was randomly chosen to be between 0 and 359, and heading angle was not updated based on wall proximity. In the second experiment, simulated flies followed momentum-retaining movement rules as described above, but did not update their heading angle depending on wall proximity. In the third experiment, we simulated flies using movement rules and wall proximity-based heading angle updates as described above. Additionally, to investigate the effect of wall-following on simulated flies’ motion and decision-making predictability readout, we performed a parameter sweep on the wall proximity-dependent angle weight averaging parameter *WF*. Over a second set of 5 additional experiments, we systematically increased *WF* from 0, which equals the previously described second experiment, to 0.1, which equals the previously described third experiment. We simulated 100 flies per experiment, chosen to approximate behavioral assay sample sizes. Flies were simulated for 108000 steps, mirroring a sampling rate of 30hz over a 1h experiment. To alleviate Brownian-motion dependent relatively low number of scorable trials, flies in the first experiment were instead simulated for 540000 steps. Resulting datasets were analyzed in the same fashion as datasets originating from behavioral assays. We analyzed movement only after reaching the lowest point in the cul-de-sac (*y*′ =0), as Brownian agents repeatedly traversed the same *y*-bin, rendering the distinction between −*y*′_*Tr,k*_ and *y*′_*Tr,k*_ meaningless.

First, we found that TPI curves of Brownian agents appeared very different from real flies (Fig. S13A). Given the large degree of noise in this ABM version, the Brownian agent displays numerous downward/upwards traversals between the bottom arm and the intersection in a single trial before eventually entering either the left or right arm (Fig. S14). This is in contrast to real flies that typically make single downward and upward traverses in each trial. Critically, with the last midline-crossing and, hence, the increase in predictability occurring only very close to the intersection, the behavior of Brownian agents substantially deviates from the behavior of real flies (c.f., Figs. S13A & 4A). Moreover, the number of midline-crossings during upward motion made by these simulated agents is many folds larger than that of flies (c.f., Figs. S6C, right & S13E, top right). Thus, the Brownian ABM does not replicate the core determinants of the motion choice dynamics observed in real flies.

Second, the behavior of *WF* =0 and Brownian agents differ in two main aspects. The average number of midline-crossings after the cul-de-sac is of a similar magnitude to those of real flies (c.f., Figs S13E, left, dark red & S6C, right). In contrast to Brownian agents and real flies, the predictability of no wall-following agents decreases *below* zero during their upward motion through the arm (Fig. S13A, top, dark red), capturing their tendency to follow the wall *incongruent* with their eventual turn direction (Fig. S15D). Increasing *WF* above zero relaxes this turn-incongruent wall-following effect (e.g., see *WF* =0.01, orange curve in Fig. S13A, top) whereas an opposite, turn-*congruent* wall-following effect is displayed by agents with sufficiently large *WF* (*WF* ≥0.05 in Fig. S13A). As *WF* is further increased, the central tendency of the distribution of last midline-crossings shifts towards lower y values (Fig. S13A, bottom), reflecting an earlier onset of wall-following (Fig. S15D) and, consequently, an earlier increase in predictability above zero (Fig. S13A, top).

Third, considering global lateral tendencies and, as could be expected from the earlier onset of their effective wall-following (Fig. S15D), agents with larger *WF* not only display TPI curves above those with lower *WF* (Fig. S13A, top) but also exhibit increased MAD scores during upward motion (Fig. S13B). This is in line with differences between WT flies and flies of most mutant lines (See: Figs. 4A, top & 4B, top). However, in contrast to real flies, *P*_*Even*_, the local tendency to make even amounts of midline-crossings after the cul-de-sac, deviates from chance only for agent with large *WF* (Fig. S13C-D: *WF* =0.09 *& WF* =0.1, dark blue & purple, respectively). That is, because, with their (effective) later onset of wall-following, the horizontal location *within* the cul-de-sac for agents with lower *WF* is almost uninformative of their eventual turn-direction (see *WF* =0.05,0.07 in Figs. S15D & S13A). Conversely, for agents with larger *WF* (*WF* =0.09,0.1), the horizontal location within the cul-de-sac is informative of the eventual turn direction. Naively, this could suggest that the way these agents make turns is somehow similar to the turn-decision process of real flies. However, upon closer examination, it becomes clear that this is not the case. This is because, in contrast to the behavior of real flies, for which predictability typically starts deviating from zero *within* the cul-de-sac (Figs 4A & S6A, middle) and is sustained by means of local lateral tendencies, a change in predictability within the cul-de-sac for all ABM versions is either absent or reflects the magnitude of predictability already when entering the cul-de-sac (*WF* ≤ 0.07 and *WF* ≥0.09, respectively, in Figs. S13A, top & S15A, center). Indeed, in more than 90% of the trials, *WF* =0.09 agents make a single midline-crossing within the cul-de-sac (Fig. S13E, right), which expresses their enduring wall-following tendency even before the cul-de-sac. This tendency is even further pronounced in *WF* =0.1 agents, which wall-follow the surrounding wall over almost the entire maze (Figs. S13A, top, & S13E). Thus, the midline-crossings made by ABM agents typically reflect their movement *before* committing to the wall to which they persist following (*WF* ≤ 0.07) or simply the endurance of wall-following that started before the cul-de-sac by means of a single midline-crossing (*WF* ≥0.09). Post wall-commitment, movement of ABM agents is almost deterministic: they typically make either zero or a single midline-crossing and therefore, a local lateral tendency metric of the parity in the amount of midline-crossing after this location adds no information about the eventual turn-direction.

Thus, the wall-following agents introduced by the ABM framework cannot replicate the capacity of real flies to sustain their probabilistic motion signatures by means of local tendencies. As TPI is tightly linked with the tendency to make even midline-crossings throughout the remainder of the trajectory, we conclude that TPI curves of real flies and simulated ABM agents reflect very different decision processes. These results further emphasize that flies do not solely rely on wall-following. That is, there must exist decision processes beyond simple movement rules that flies rely on when making decisions across space and time.

### Quiver plots

To create the average quiver plots shown in Fig. 4D, we computed two primary variables for each fly in each genetic line under four conditions: left turns (blue), right turns (red), and motion towards (*y* < 0) and away *y* >0) from the cul-de-sac edge. These variables were the spatial density and the average movement direction. In what follows, we describe these computations for an individual fly and finally, how each average quiver is computed across flies.

The cul-de-sac was divided into a 28-by-22 2D grid with bins defined by *X* = {−0.2268, …, 0,0.0162, …, 0.2268} and *Y* = {0,0.0162, …, 0.3564} (or *Y* = {−0.3564, −0.0162, …, 0} for *y* < 0). For each frame *f* of a fly’s trajectory within the cul-de-sac, we determined the bin (*i, j*) corresponding to the coordinates (*x*_*f*_,*y*_*f*_) and incremented the count for that bin: 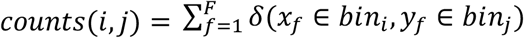, where *δ* is the indicator function that equals 1 if (*x*_*f*_,*y*_*f*_) is within bin (*i, j*) and 0 otherwise, and *F* is the total number of frames across all trials. To compute the average movement direction for each frame *f*, we calculated the velocity vector *ν*_*f*_ = (*x*_*f*+1_ − *x*_*f*_,*y*_*f*+1_ −*y*_*f*_) and computed the corresponding angle *θ*_*f*_ = *arctan*2(*y*_*f*+1_ −*y*_*f*_, *x*_*f*+1_ − *x*_*f*_). For each bin (*i, j*), we accumulated these angles and their corresponding magnitudes |*ν*_*f*_|. The magnitude-weighted average angle *θ*_*i,j*_ for bin (*i, j*) was computed by converting each angle *θ*_*f*_ to its complex representation 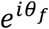 and computing the weighted sum of these complex representations: 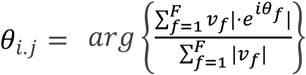

To compute the average quiver over multiple flies, we combined the corresponding *counts*_*i,j*_ and *θ*_*i,j*_ from individual flies. The *counts* of each fly were normalized to represent the relative frequency of visits to each bin, *freqs*_*i,j,n*_, where *n* indexes the flies. The average direction angles (*θ*_*i,j*,)_) for each fly were converted to their complex representation,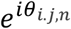. For each bin, we computed a frequency-based weighted average of the angles across all flies: 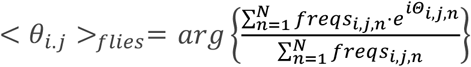. The relative frequency of visits to each bin was averaged over all flies to get the average frequency with which each bin was visited: 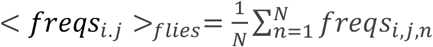. The computed average direction (< *θ*_*i,j*_ >_*flies*_) and frequency (< *freqs*_*i,j*_ >_*flies*_) for each bin were used to plot the average quiver. The direction of each vector in the average quiver plot corresponds to the average direction of movement, while the magnitude of each vector represents the relative frequency of visits to that bin.

### Heat maps

In addition to the quiver plots, we computed average heat maps over flies (or simulated agents) to visualize the spatial distribution of their locations during motion away from the cul-de-sac *y* >0. In what follows, we describe the computation for an individual fly and finally, how the average heatmap is computed across flies.

The maze was divided into a 2D grid of bins defined by *X* = {−0.1944, …, 0,0.0324, …, 0.1944} and *Y* = {0,0.0324, …, 1.4580,1.4904}. The spatial probability for each bin (*i, j*) was defined by: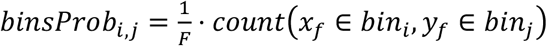, where *f* is a frame in the trajectory data of a fly, *F* is the total number of frames for that fly that lay in the 2D grid and *count* is the number of frames where (*x*_*f*_,*y*_*f*_) falls within bin (*i, j*).

To compute the average heat map over multiple flies for each turn direction (left or right), the spatial probability (*binsProb*) for each fly was computed separately for left and right turns, and then normalized to represent the probability distribution across bins (Fig. 2B, top and center). For each bin (*i, j*), the average probability was computed across all flies: 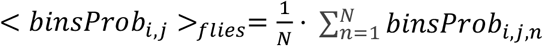. To visualize the differences between left and right turns, we also computed 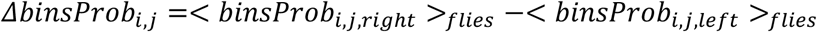 (Figs. 2B, bottom, & S15D).

### Zero-midline-crossings comparison between mutants and WT

To assess the contribution of trials with zero midline-crossings to the sharper increase in predictability within the cul-de-sac for WT and mutant flies, we computed *TRI*(*yRange*) after excluding trials in which there were zero midline-crossings after the cul-de-sac. While this exclusion involves an artificial debiasing of the parity tendencies of all flies and thus expected to result in reduced predictability (i.e., because excluding zero midline-crossing trials necessarily reduces the probability to make turns congruent with the horizontal location in the cul-de-sac towards chance level), we use it as a metric to estimate the departure in predictability between WT and mutant flies with respect to their horizontal location just before they leave the cul-de-sac. Figures S6D and S6A, bottom depict the average TPIs for positive midline-crossings and their corresponding estimates for the change in predictability within the cul-de-sac. While FoxP, NorpA, and NompC mutant lines all differed significantly in their cul-de-sac TPI values when considering all trials, a substantial effect persisted primarily in Dumb flies after zero-omission (Fig. S6A, top).

### Simulated midline-crossings

To simulate the distributions of #*crossings* for mazes of different arm length we considered the midline-crossings made by flies in the long maze. Recall that #*crossings* is defined as the number of midline-crossings during upward motion away from the cul-de-sac (Fig S5A; see also Fig. 3D). Because flies in the long maze rarely make midline-crossing after leaving the intersection (0.011%, 4 crossings out of 35,759 post cul-de-sac midline-crossings made by flies in the long maze) then effectively, these midline-crossings typically occur in the interval between the upper edge of the cul-de-sac and the upper edge of the intersection (see Fig. S3B-C). Specifically, the length of this interval is 2*d* + *k* in the long maze, where 2*d* is the length of each arm in the long maze and *k* is the length of the intersection (between its bottom and top ends). The corresponding length of this interval in the short maze is *d* + *k* and, for any maze with different arms’ length, the interval can be similarly described as *rd* + *k*, where *r* is the arm-length ratio of any maze to the short maze. To consider the number of midline-crossings expected in mazes with different arm length (*r* = {.1, .33, .5, .66, .9, 1, 1.1, 1.33, 1.5, 1.66, 1.9, 2}) we computed, for each trial made by flies in the long maze, the number of (the first) midline-crossings occurring before the *rd* + *k* interval is exceeded. Bootstrapping trials from the resulting trimmed #*crossings* for each *r* results in the expected distributions seen in Figure S5D. The resulting #*crossings* distribution for *r* = 1 replicates the observed #*crossings* distribution of flies in the short maze and hence the tendency to make even #*crossings* across the sample. This tendency is also expected from mazes with arm-length either larger (*r* > 1) or smaller than the short maze (*r* < 1).

We also simulated the #*crossing* distributions expected in the short and long mazes under the assumption that midline-crossings are given by draws from an inter-crossing-interval distribution. For this purpose, we computed for each fly and each trial in the long maze the inter-crossing intervals. For each trial, the y-locations of all post cul-de-sac midline-crossings are given by *y*_*j*_|*cross*, where 1 ≤ *j* ≤ *k* and *k* is the number of post cul-de-sac crossings in that trial. Defining *y*_0_|*cross* and *y*_*k*+1_|*cross* as the y-values denoting the y-locations of the cul-de-sac upper edge and the intersection upper edge (respectively; similar across all flies and trials), the inter-crossing-distances in that interval, measured in distance only on the y-axis, is given by *Δy*_*j*_ =*y*_*j*_|*cross* −*y*_*j*−1_|*cross*, where 1 ≤ *j* ≤ *k* + 1. To consider the expected distribution of #*crossings* under the above assumption in either the short or long mazes, we sampled #*trials*(*f*) trials from each fly in the ling maze, where #*trials*(*f*) is the number of trials made by fly *f* in the long maze. For each simulated trial, we sampled *Δy* values from all *Δy* across all trials made by this fly. To avoid over-sampling of trials with numerous #*crossings* (and hence small *Δy*’s) the sampling weight (probability) 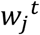 of each 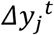,an inter-crossing-distance of the fly in trial *t*, was set to [(*k* + 1) · #*trials*(*f*)]^−1^. For each trial, *Δy*s were sampled until 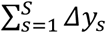,the accumulated *Δy*’s in the simulated trial reached the interval between the upper edge of the cul-de-sac and the upper edge of the intersection (long maze: 2*d* + *k*, short maze: 2*d* + *k*). The resulting number of midline-crossings in the simulated trials is given by the *S* − 1, where *S* is the number of sampled *Δy*’s when the interval is crossed for the first time. Simulating #*trials*(*f*) for each fly *f* for each of the flies resulted in one simulated distribution of #*crossings* under the above assumption of draws from a distribution of inter-crossing-distances. Repeating this procedure 1000 times results in the expected distributions in Fig. S5B. These simulations did not replicate the observed distributions of #*crossings* in short and long mazes (Fig. S5B). In an attempt to test whether this relates to some differences between midline-crossing made within the maze’s bottom arm and those made within the intersection, we repeated the above process by computing and simulating only *Δy* and 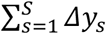 occurring within the arm (intervals for long and short mazes: 2*d* and *d*, resp.). The simulated #*crossings* in a trial of that revised procedure was given by the sum of *S* − 1, the number of sampled *Δy*’s when the revised interval (2*d* or *d*) was crossed for the first time and the observed number of midline-crossings occurring within the intersection in that trial. This revised procedure failed to replicate the observed distributions of #*crossings* in short and long mazes (Fig. S5C).

**Table S1.**
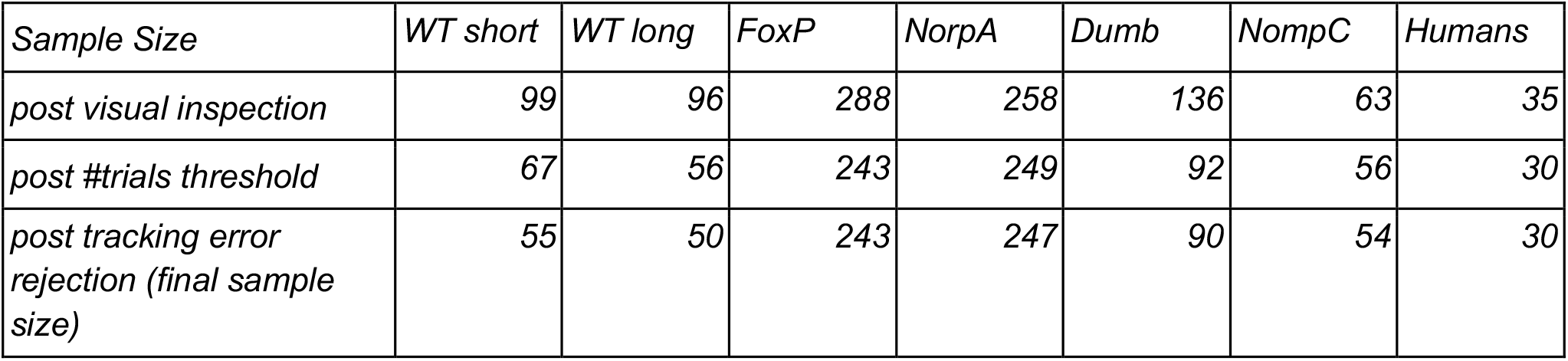
Sample sizes and exclusion criteria.

**Figure S1.**
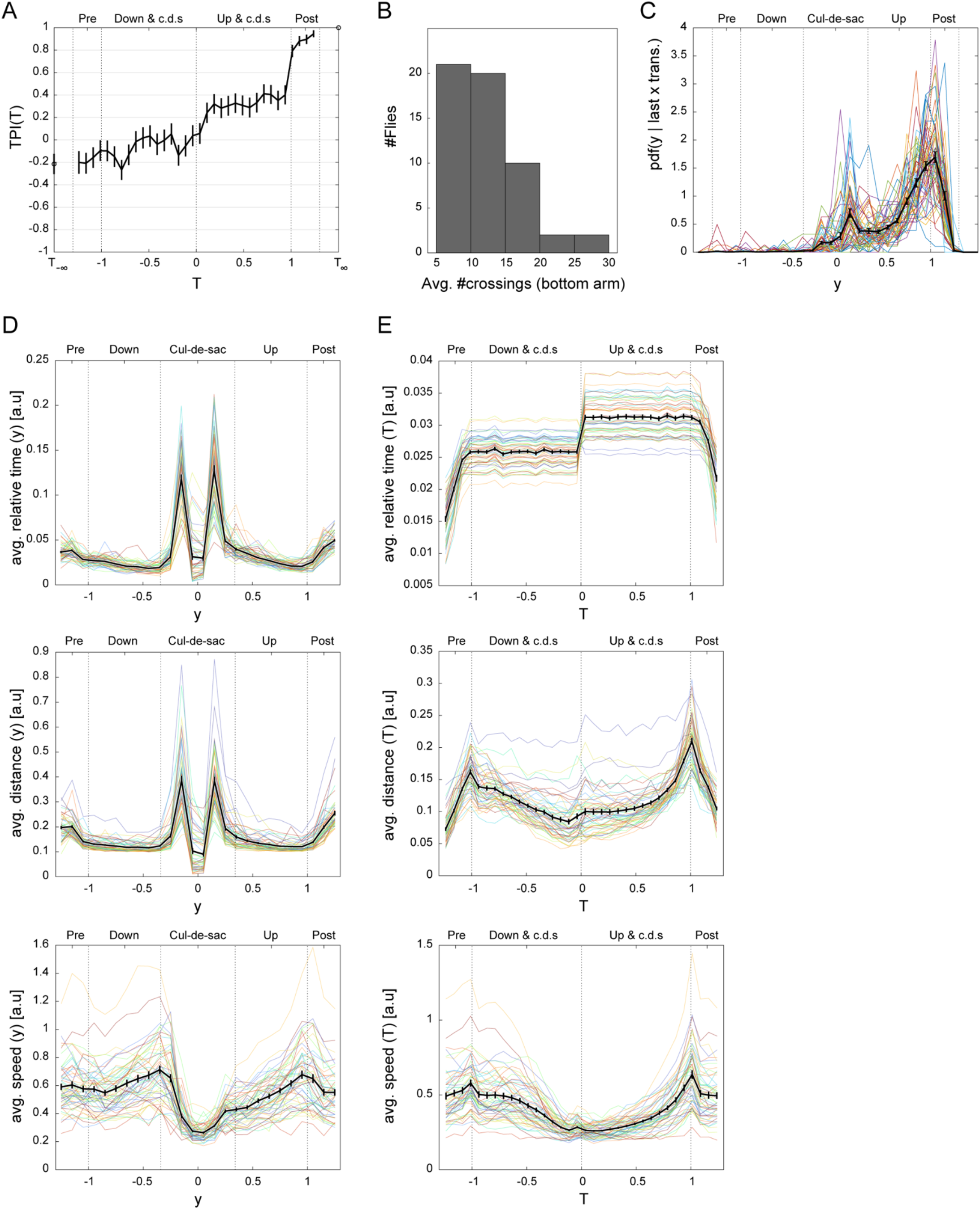
Flies in the short Y-Maze. (A) Turn Predictiveness Index (TPI) in the temporal domain for the fly in Fig. 1B-D. *TRI*(*tRange*) measures, for a given *tRange*, how predictable the average x location of a fly is with respect to its forthcoming turn direction (see Materials and Methods). Error bars denote Standard Error (SE). (B) Average midline-crossing histogram. The histogram denotes the trial-average amount of midline-crossings (passes through *x* =0) made by each fly (*n* =55) within the bottom arm in each trial (−1 < *y* < 1). (C) Average PDF of last midline-crossings (LMC). LMC is defined as the last *y*-location within a given trial in which the *x*-location changed polarity (i.e., last crossing of the horizontal midline). The average *PDF* (*y* |*LMC*) curve (Black) over the entire sample (*n* =55; WT) is overlaid on the *PDF*(*y* |*LMC*) curves of individual flies. Error bars denote Standard Error of the Mean (SEM). (D) Average motion kinematics of flies across the maze, depicting average relative time (top), distance (center) and speed (bottom), computed for the same *y*-bins in Fig. 1F. The average curves across flies (Black; *n* =55) is overlaid on the trials-average curves of individual flies. (E) As in D for the *relative* time-bins in Fig. 1G.

**Figure S2.**
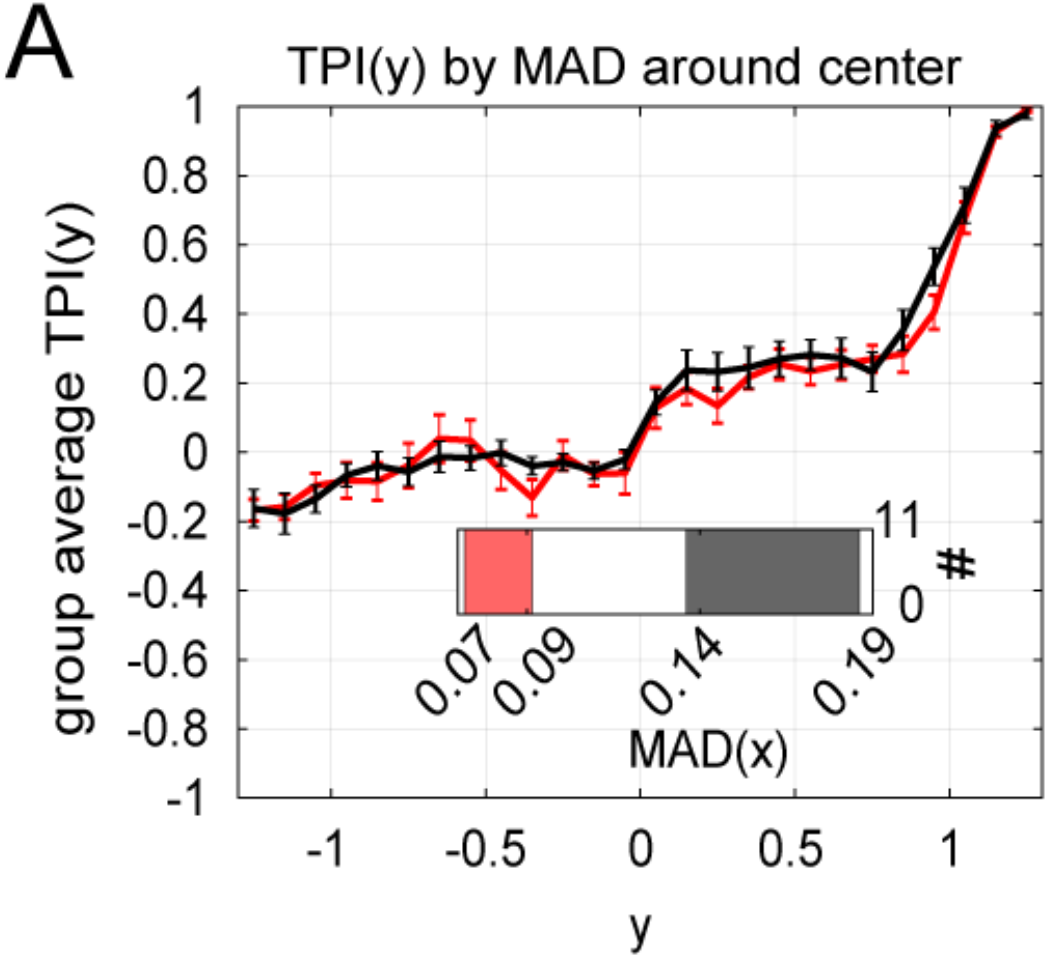
Average TPIs for flies in the short maze with lowest (red) and highest (nlack) *MAD*(*x*| − 1 < *y* < −0.34) (*n* = 11 in each group; WT). Inset: distribution of *MAD*(*x*| − 1 < *y* < −0.34) values over the sample. The *MAD*(*x*| − 1 < *y* < −0.34) value of each fly computes the median absolute deviation from the horizontal midline for *downward* motion from the upper edge of the bottom arm until entering the cul-de-sac (c.f., Figs. S2 & 2E).

**Figure S3.**
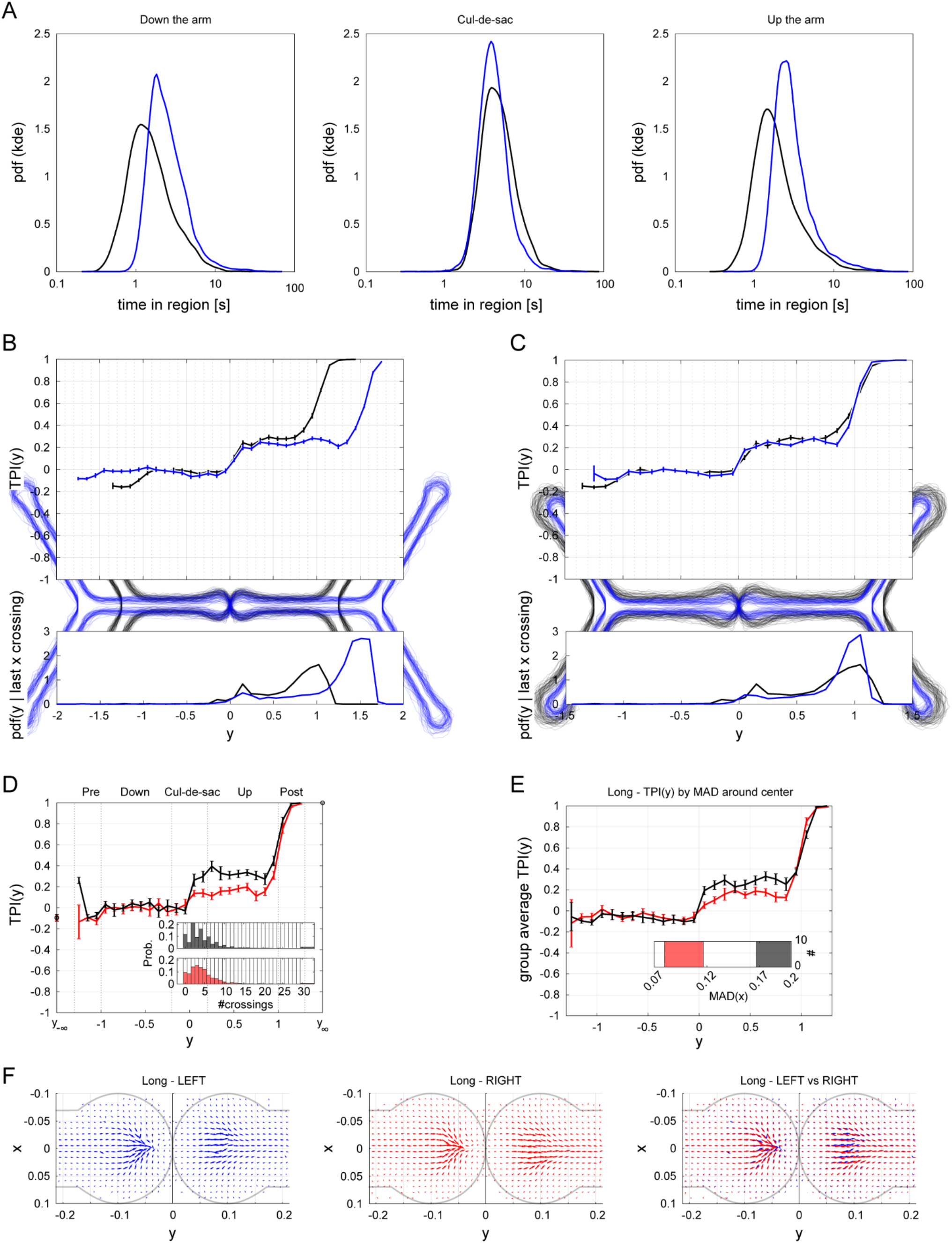
Long vs short mazes in the spatial domain. (A) Probability density functions (PDFs) of the absolute (to not confuse with relative) time spent walking down the arm (left), within the cul-de-sac (middle) and walking up the arm (right) for WT flies in the short (black; *n* =55) and long (blue; *n* = 50) mazes. Each curve depicts a Kernel smoothing function estimate over log10 of the time spent in all trials across the sample. (B) Average TPI curves (top) and corresponding PDF of last midline-crossings (bottom) in absolute-size representation. All bins are of equal width and represent equal size segmentations of the raw plane (background images). (C) As in B, for relative representation of the plane, with *y* = ±1 defining the upper edge of the bottom arm (bottom edge of the intersection). (D) Average TPIs for flies in the long maze with lowest (red) and highest (black) *P*_*Even*_ values (calculated as in Fig. 3D, Top, blue; n=10 in each group). Inset: Probability mass functions of #*crossings* during upward motion away from the cul-de-sac in each group (color coded as in the main panel; #*crossings* def. as in Fig. 3D, bottom). (E) As in D, for global lateral tendencies in the long maze: Average TPIs for flies with lowest (red) and highest (black) MAD scores (*n* = 11 in each group). Inset: distribution of (MAD scores during upward movement through the arm) over the sample. (F) Quiver plots depicting average motion within the cul-de-sac across flies in the long maze. Y-coordinates in D-F are standardized as in C. For long vs short: c.f., Figs. S3D & 2D, S3E & 2E, S3F & 2A.

**Figure S4.**
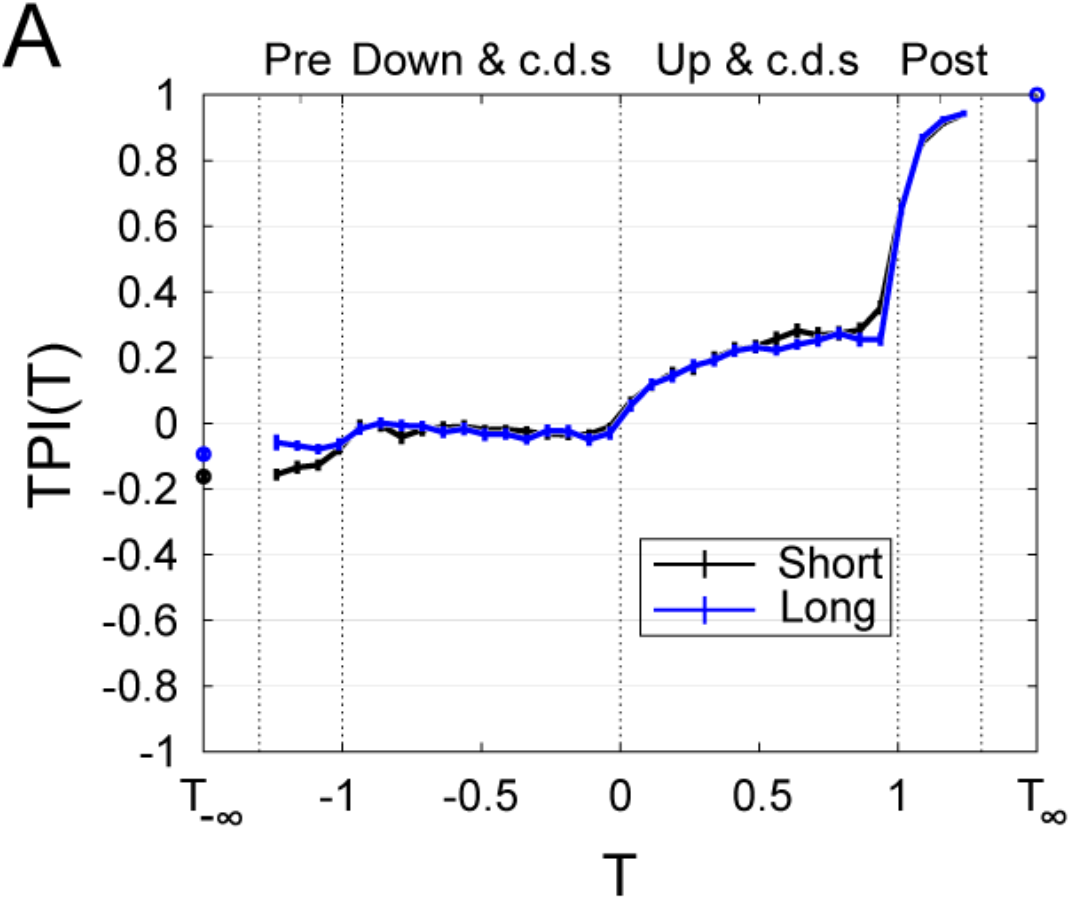
Long (blue; *n* = 50) vs short (black; *n* =55) average TPI curves in the temporal domain. Error bars denote Standard Error of the Mean (SEM).

**Figure S5.**
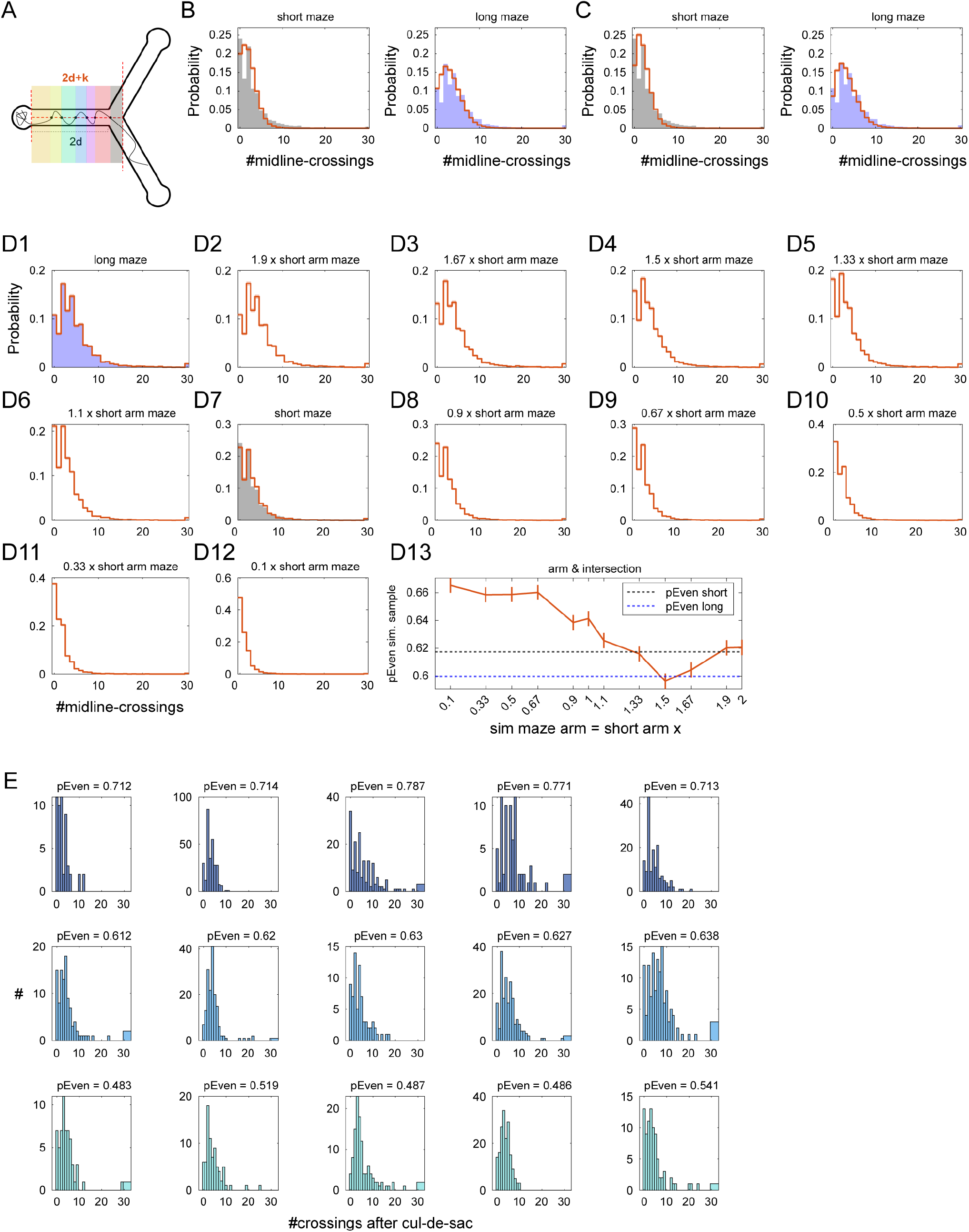
Post cul-de-sac midline-crossings in the long maze and corresponding crossing simulations. (A) Illustration of post cul-de-sac midline-crossings in a trial. After leaving the cul-de-sac, the illustrated trajectory passes through the horizontal midline 6 times and, thus, #*crossings* = 6. The same definition for #*crossings* is used in Figs. 2C-D, 3D, 4C, S3D, S6C, right, S6D, S9, S10, S11B, and S13C-E, left. Because flies in the long maze almost never make midline-crossing after the intersection (0.011%, 4 crossings out of the 35,759 post cul-de-sac midline-crossings in Fig. 3D, bottom, blue) then, effectively, post cul-de-sac midline-crossings are constrained within a 2*d* + *k* interval between the upper edge of the cul-de-sac and the upper edge of the intersection, where 2*d* is the length of the arm in the long maze (*d* in the short maze, *r* · *d* in any maze with arms’ length *r* times the short maze) and *k* is the length of the intersection (similar across mazes). (B) Simulating inter-crossing-y-intervals does not replicate the observed #*crossings* distributions in the short (left) and long (right) mazes. Filled histogram: observed #*crossings* pdf (as in Fig. 3D, bottom). Orange line and shaded area: simulated #*crossings* pdf, averaged over 1000 simulations and simulations’ std, respectively. In each simulation (*n*_*sims*_ = 1,000) and each fly in the long maze (*n* = 50) we simulated the number of crossings in each trial by sampling from the fly’s observed post cul-de-sac inter-crossing-y-distances across all trials (see Supporting Information). The simulated number of post cul-de-sac crossings in each trial was determined when the accumulated inter-crossing-y-distances in a simulated trial attained *d* + *k* or 2*d* + *k* for the short (left) and long (right) maze simulations, respectively. (C) As in B, simulating the number of post cul-de-sac crossings in the short (left) and long (right) mazes. But, using the inter-crossing-y-intervals that occur only *within* the arm of the long maze (in a 2*d* interval) to simulate the number of midline-crossings within the arm (short maze: *d* + *k*, long maze: 2*d* + *k*) and adding the observed number of crossings in the intersection (*k*) as the overall simulated number of post cul-de-sac crossings in a trial. (D) Bootstrapping the (trimmed) #*crossings* made in the long maze preserves parity crossings tendencies in shorter mazes and replicates the observed #*crossings* distribution in the short maze. To consider the expected distributions of #*crossings* we counted, for each trial in the long maze, the number of first crossing attained within the interval *r* · *d* + *k*, where *r* = {2, 1.9, 1.66, 1.5, 1.33, 1.1, 1,0.9,0.66,0.5,0.33,0.1} (D1-D12, respectively). We computed the expected distributions for each maze with arm length *r* by bootstrapping trials from the corresponding trimmed post cul-de-sac #*crossings* (*n*_*samples*_ = 10,000 and *n*_*trials*_= 7,360 in each sample). Orange line and shaded area: expected #*crossings* pdf in the arm and intersection (a. & i.), averaged over 10,000 simulations and the simulations’ std, respectively. Filled histogram in D1 & D7: observed #*crossings* pdf (as in B, C and Fig. 3D, bottom). D13: Fraction of even post cul-de-sac #*crossings* across all trials in the sample. Dashed: fraction of even #*crossings* observed in the short (black) and long (blue) mazes. Orange: average (line) and std (error bars) of the fraction of even #*crossings* across samples for each *r*. (E) Distributions of post cul-de-sac #*crossings* of individual flies in the long maze with largest (top) intermediate (center) and smallest (bottom) *P*_*Even*_ values (real data).

**Figure S6.**
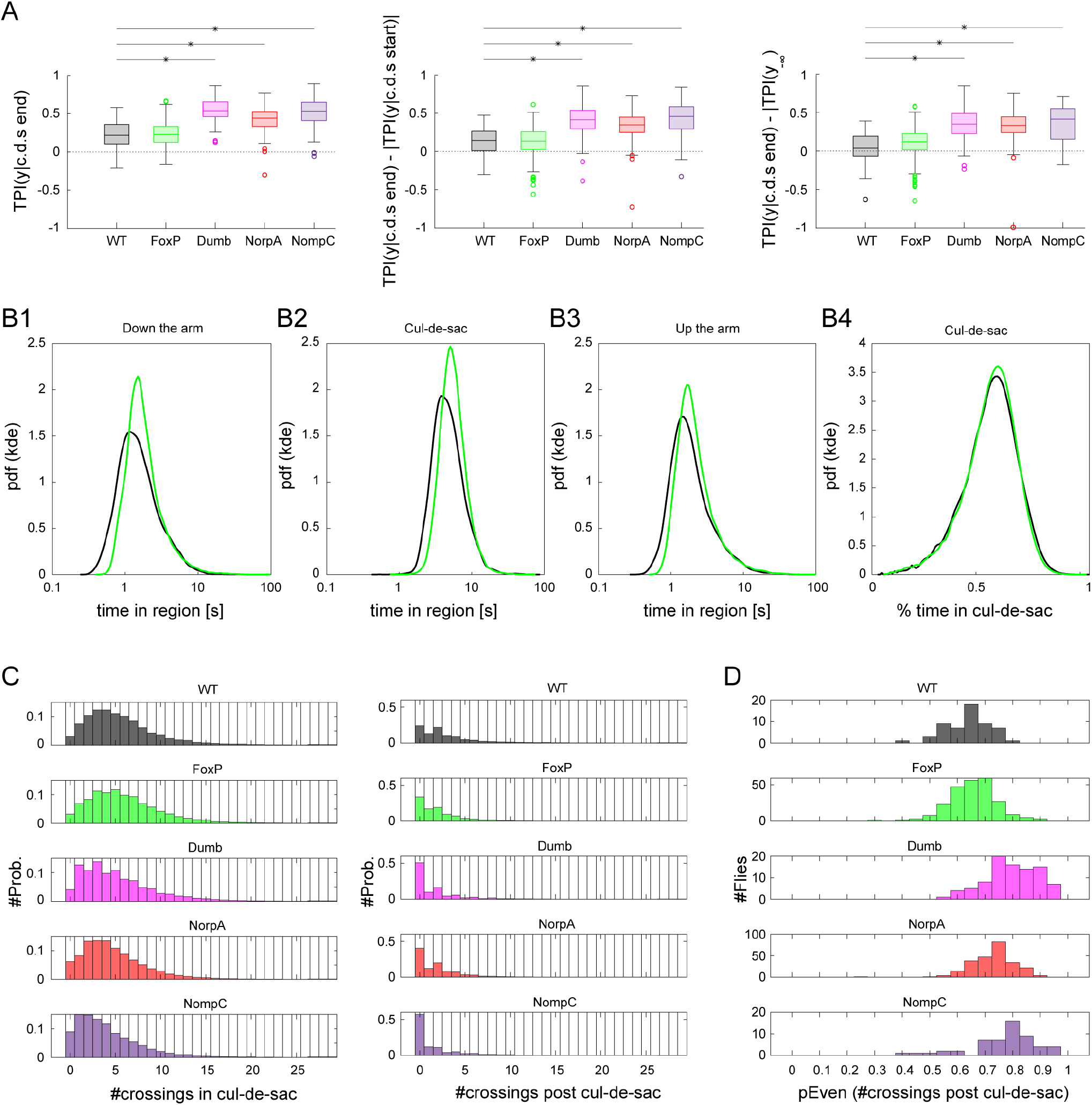
Mutants and WT comparisons. (A) TPI increase in the cul-de-sac across genetic lines: WT (black, *n* =55), FoxP (green, *n* = 243), dumb (magenta, *n* = 90), NorpA (red, *n* = 247) and NompC (purple, *n* = 54). For each fly in each line, we computed the TPI value, *TRI*(*yRange*), at the last location within the cul-de-sac (*yRange* = {0.2 ≤ *y* ≤ 0.3}, left), its difference with the TPI value at the last location within the cul-de-sac (*yRange* = {−0.3 ≤ *y* ≤ −0.2}, middle) and its difference with the left tail value of the *TRI*(*Y*_−∞_) (right). Each box plot depicts median (central mark), 25th and 75th percentiles (bottom and top edges of the box), most extreme data points not considered outliers (whiskers), and outliers (individual circles) across a genetic line. Asterisks depict significant differences between WT and mutant line (two-sided Wilcoxon rank sum test, corrected for multiple comparisons: *α* =0.05/4; left: *p*_*WT,FoxP*_ =0.67, *p*_*WT,FoxP*_ < 0.001, *p*_*WT,dump*_ < 0.001, *p*_*WT,NorpA*_ < 0.001, middle: *p*_*WT,FoxP*_ =0.87, *p*_*WT,FoxP*_ < 0.001, *p*_*WT,dump*_ < 0.001, *p*_*WT,NorpA*_ < 0.001, right: *p*_*WT,FoxP*_ =0.07, *p*_*WT,FoxP*_ < 0.001, *p*_*WT,dump*_ < 0.001, *p*_*WT,NorpA*_ < 0.001). (B) Probability density functions (PDFs) of the absolute (to not confuse with relative) time spent walking down the arm (B1, −1 < *y* < −0.34), within the cul-de-sac (B2, −0.34 < *y* < 0.34) and walking up the arm (right, −0.34 < *y* < 1) for WT (black) and FoxP (green) flies in the short. Each curve depicts a Kernel smoothing function estimate over log10 of the time spent in all trials across the sample. B4: Pdf of the fraction of time spent within the cul-de-sac (out of the total time spent between −1 < *y* < 1) across all trials in a sample, using a linear kernel smoothing function estimate. (C) Distributions of the number of midline-crossings within the cul-de-sac (left) and post cul-de-sac (right, #*crossings*) across all trials made by flies in a genetic line (color-coded as A; *n*_*trials,WT*_ = 7904, *n*_*trials,FoxP*_ = 34,764, *n*_*trials,dumb*_ = 12,391, *n*_*trials,NorpA*_ = 54,711, *n*_*trials,NompC*_= 8,255,). (D) Distributions of *P*_*Even*_, the fraction of trials with even post-cul-de-sac #*crossings* across flies in each genetic line (colors and sample-sizes as A).

**Figure S7.**
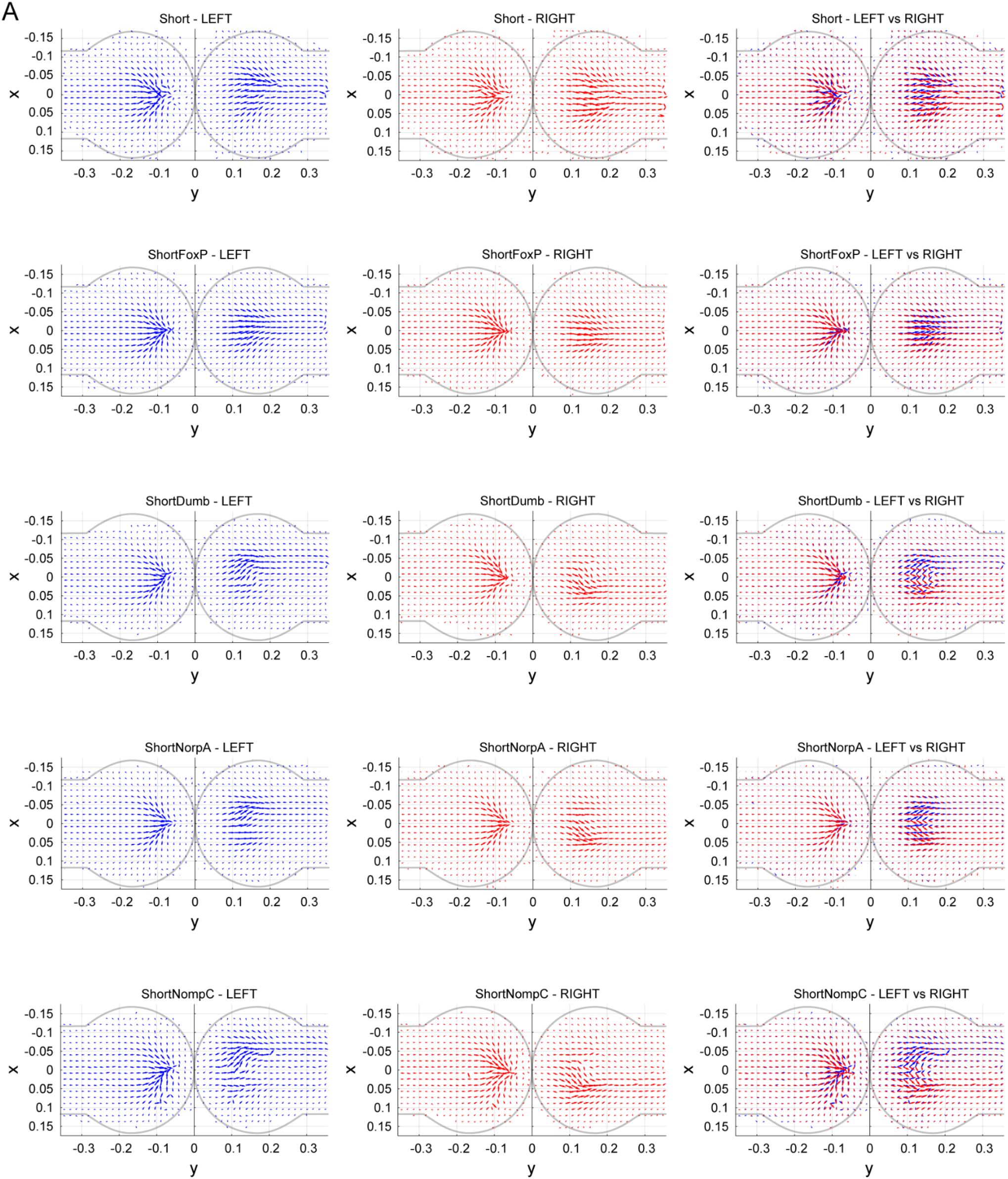
Quiver plots depicting average motion within the cul-de-sac across flies in each genetic line: WT (*n* =55), FoxP (*n* = 243), dumb (*n* = 90), NorpA (*n* = 247) and NompC (*n* = 54). Quivers are computed separately for inward (*y* < 0) and outward (*y* >0) motions and separately for left turns (left panel, blue) and right turn (middle panel, red), based on flies’ corresponding velocity vectors (Supporting Information). In each 2D bin, the vector direction represents average motion direction across all flies, while the vector length indicates the relative frequency of visits to that bin (Supporting Information). Right panel: left and right turns, overlaid.

**Figure S8.**
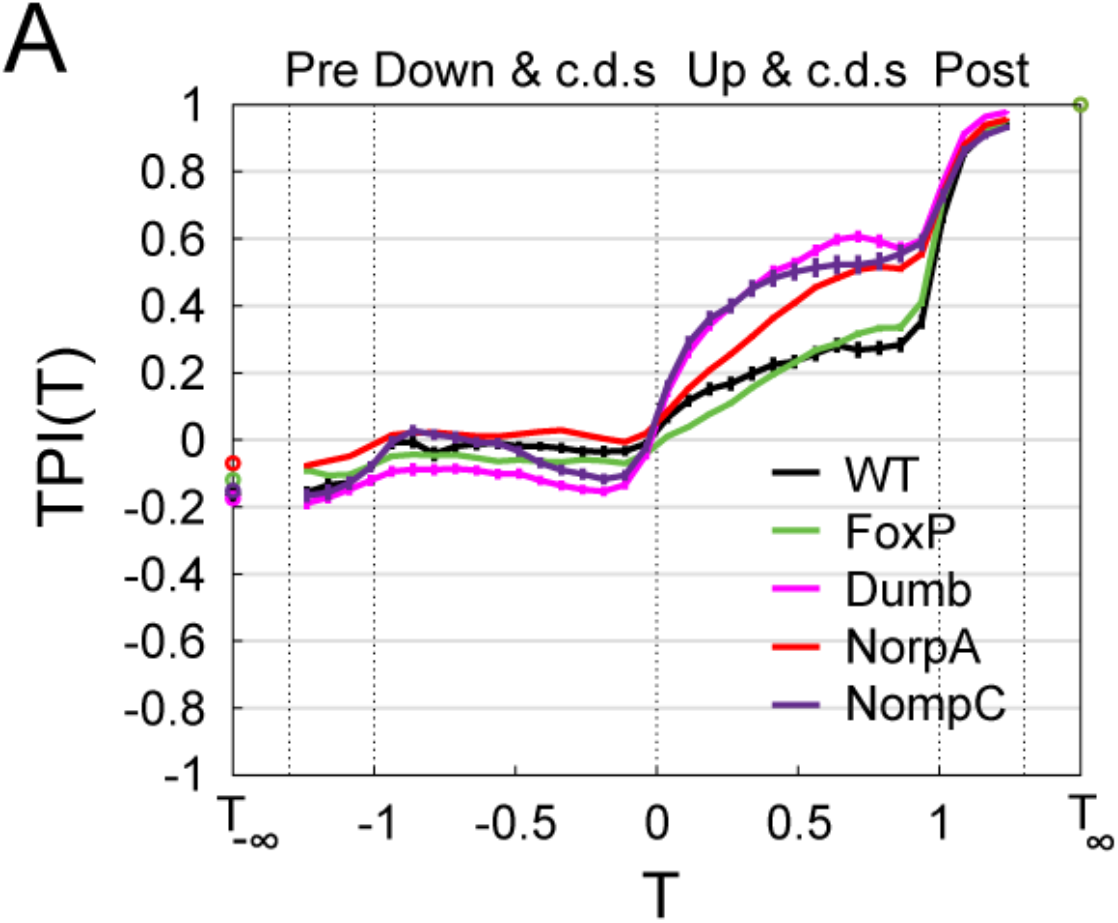
Average TPI curves in the temporal domain across genetic lines: WT (black, *n* =55), FoxP (green, *n* = 243), dumb (magenta, *n* = 90), NorpA (red, *n* = 247) and NompC (purple, *n* = 54). Error bars denote SEM.

**Figure S9.**
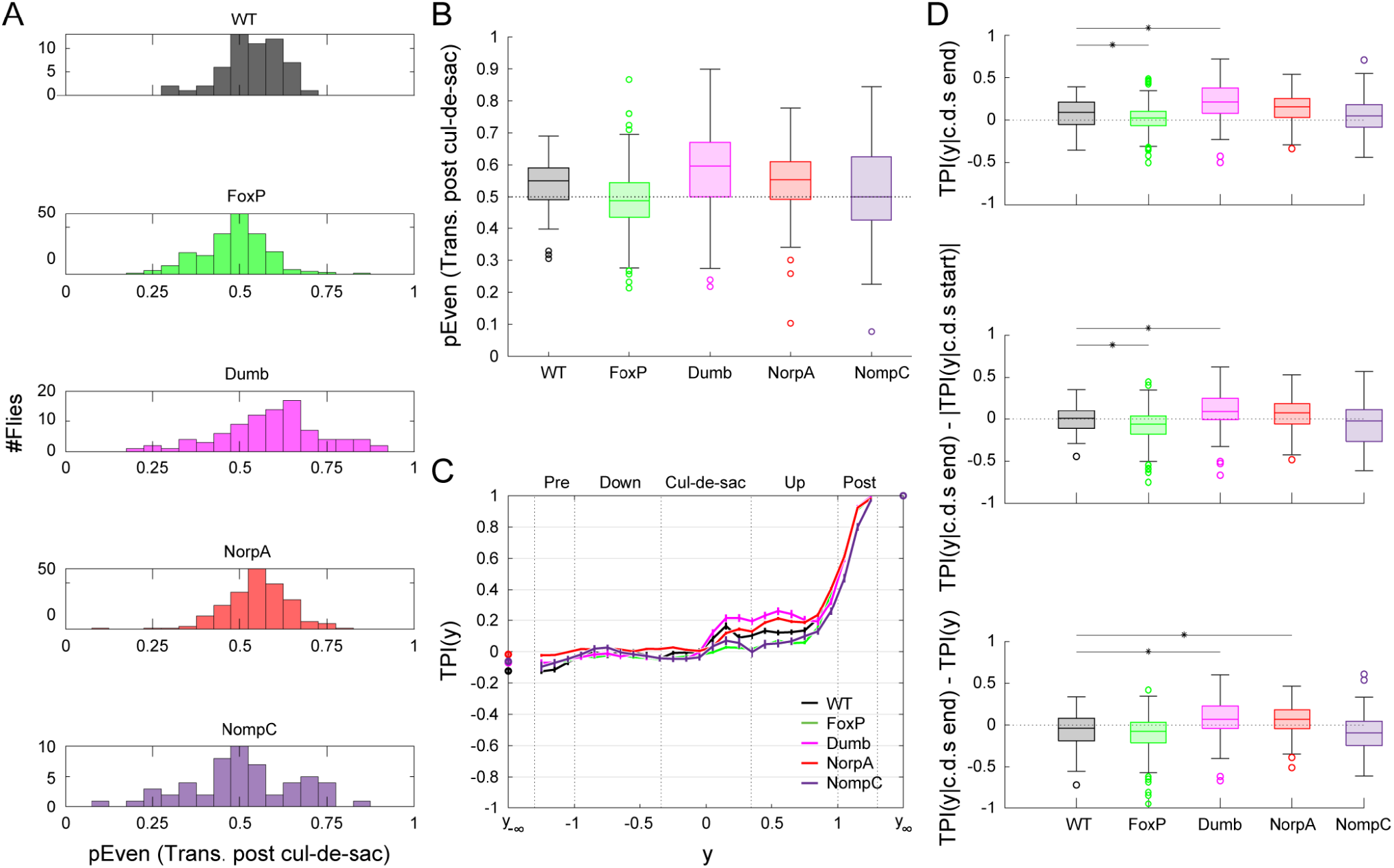
Exclusion of zero-crossings trials. (A) Distributions of the fractions of trials with even #*crossings* out of all trials with non-zero post cul-de-sac midline-crossing across flies in each genetic line: WT (*n* =55, black), FoxP (*n* = 243, green), dumb (*n* = 90, magenta), NorpA (*n* = 247, red) and NompC (*n* = 54, purple). (B) Box plots summarizing the fractions in A across flies in each generic line, color-coded as in A. Each box depicts the median (central mark), 25th and 75th percentiles (bottom and top edges of the box), most extreme data points not considered outliers (whiskers), and outliers (individual circles) across a genetic line. Across all trials in a genetic line, the fractions of even, non-zero post cul-de-sac midline-crossing are: 0.5409 ±0.0064, 0.4864 ±0.0033, 0.6025 ±0.0063, 0.5539 ±0.0028, 0.5125 ±0.0084 (*fractionon* ± *SE*) for the WT, FoxP, dumb, NorpA and NompC lines, respectively (see Fig. S10 for the fractions of trials with zero post cul-de-sac midline-crossings), and 0.5746 ±0.0061 for WT flies in the long maze (not shown). (C) Average TPI curves in the spatial domain across genetic lines, after excluding trials with zero post cul-de-sac midline-crossings, color-codes and sample sizes as in A. Error bars denote SEM. (D) TPI increases in the cul-de-sac across genetic lines (as in 6A), after excluding trials with zero post cul-de-sac midline-crossings. For each fly in each line, we excluded all trial with zero post cul-de-sac midline-crossings and computed the TPI value over the remaining trials at the last location within the cul-de-sac (*yRange* = {0.2 ≤ *y* ≤ 0.3}, top), its difference with the TPI value at the last location within the cul-de-sac (*yRange* = {−0.3 ≤ *y* ≤ −0.2}, center) and its difference with the left tail value of the *TRI*(*Y*_−∞_) (bottom). Each box plot depicts median (central mark), 25th and 75th percentiles (bottom and top edges of the box), most extreme data points not considered outliers (whiskers), and outliers (individual circles) across a genetic line. Asterisks depict significant differences between WT and mutant line (two-sided Wilcoxon rank sum test, corrected for multiple comparisons: *α* =0.05/4; top: *p*_*WT,FoxP*_ =0.004, *p*_*WT, dump*_ < 0.001, *p*_*WT, NorpA*_ =0.036, *p*_*WT, NompC*_ =0.328, center: *p*_*WT,FoxP*_ =0.002, *p*_*WT, dump*_ =0.005, *p*_*WT, NorpA*_ =0.040, *p*_*WT, NompC*_ =0.222, bottom: *p*_*WT,FoxP*_ =0.079, *p*_*WT, dump*_ < 0.001, *p*_*WT, NorpA*_ < 0.001, *p*_*WT,NompC*_ =0.234). Color-codes and sample sizes are as in A.

**Figure S10.**
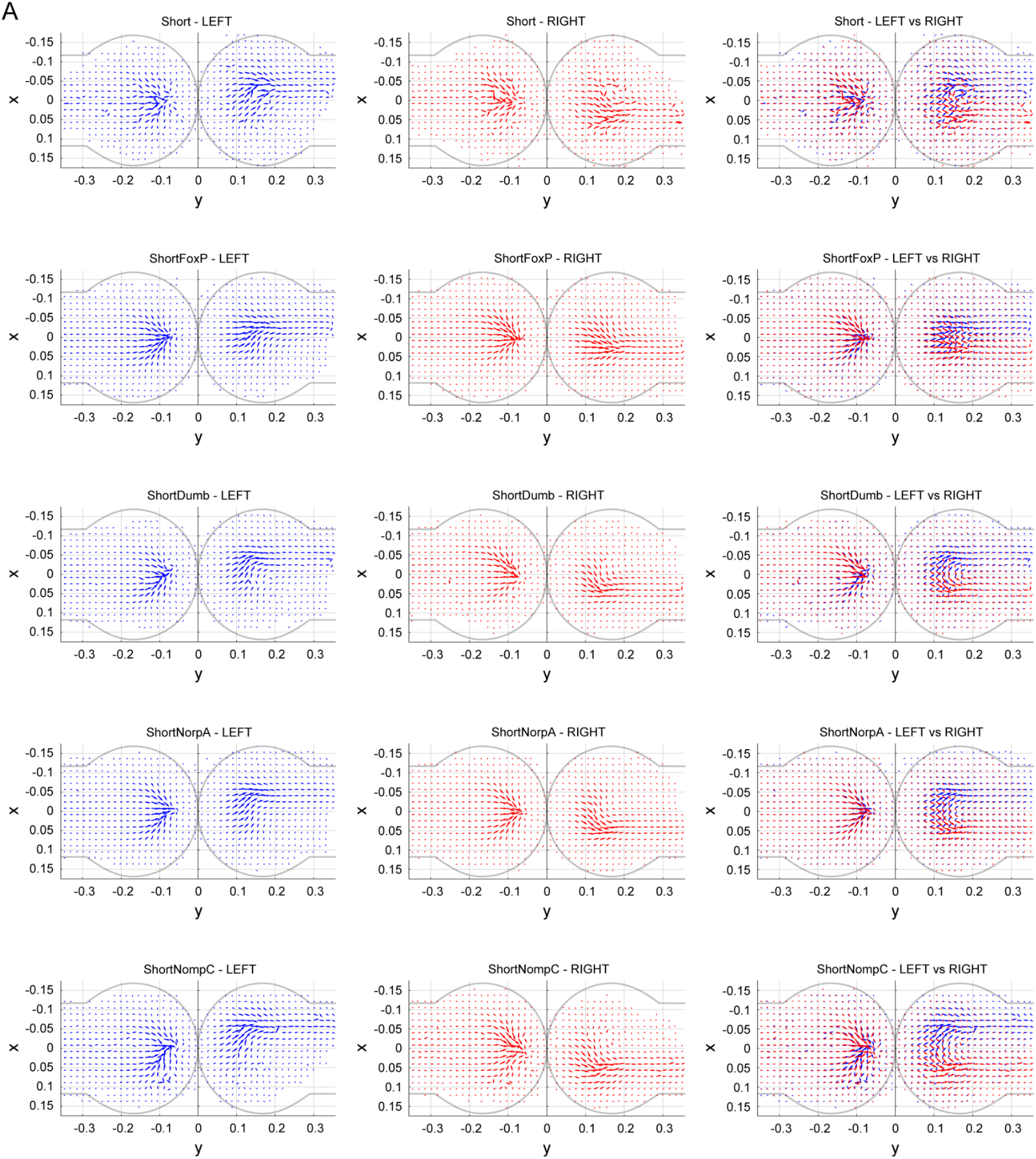
Zero-crossings quiver plots. Quiver plots as in S7, but including only trials in which there were no post cul-de-sac midline-crossings (trials in which #*crossings* =0; 24%, 34%, 51%, 24%, 40%, and 59% of the trials in the WT, FoxP, dumb, NorpA, and NompC lines, respectively).

**Figure S11.**
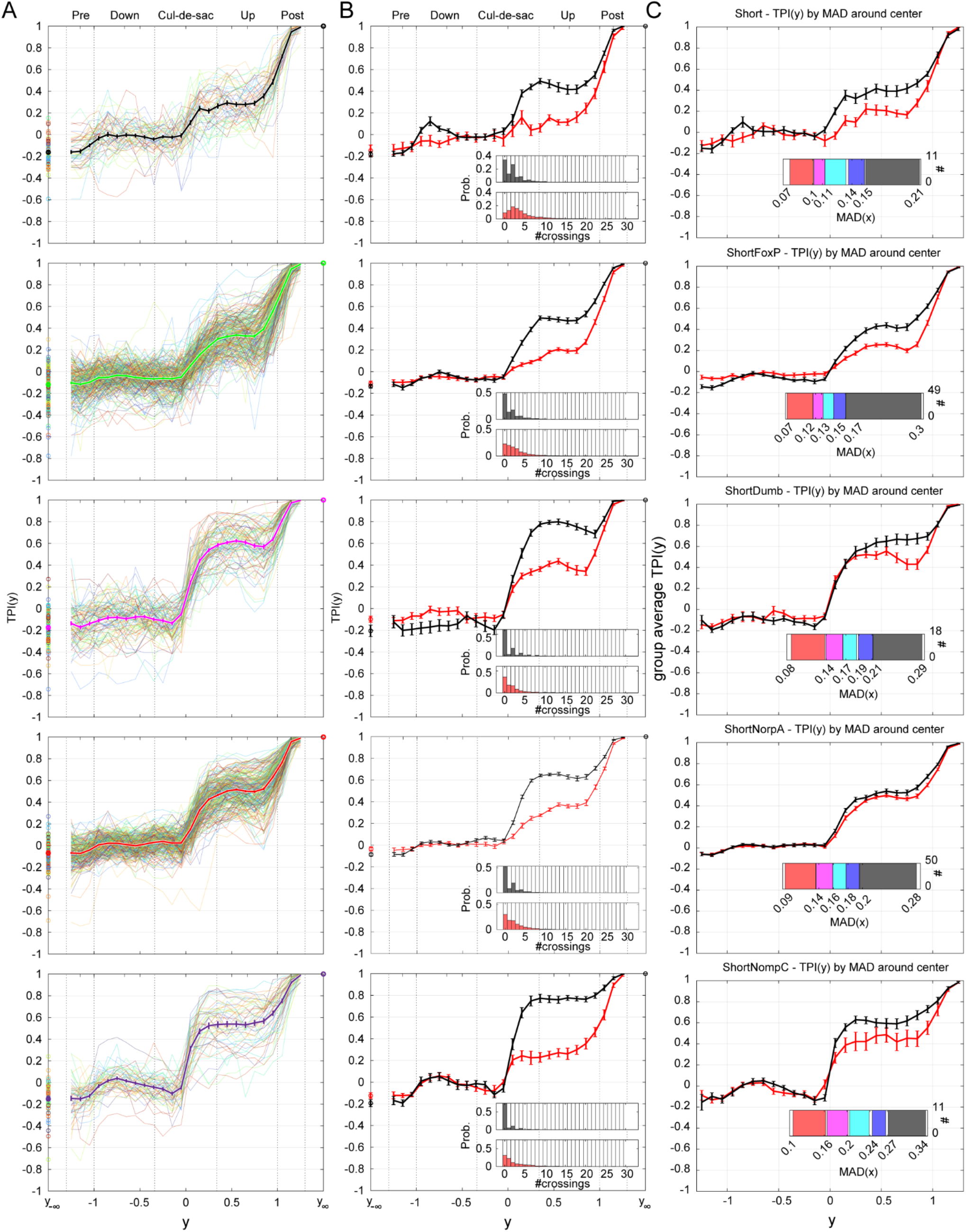
TPI of mutant and WT and their dependence on lateral tendencies. (A) Average TPI curve in the spatial domain across genetic lines (top to bottom: WT: black, *n* =55; FoxP: green, *n* = 243; dumb: magenta, *n* = 90; NorpA: red, *n* = 247; NompC: purple, *n* = 54), overlaid on the TPI curves of individual flies. Error bars denote SEM. (B) Average TPIs for flies with lowest (red, 20% of the flies) and highest (black, 20% of the flies) *P*_*Even*_ values in each genetic line (computed for each genetic line as in Fig. 2D). Inset: Probability mass functions of #*crossings* in each group (color coded as in the main panel). (C) As in B, for global lateral tendencies: Average TPIs for flies with lowest and highest MAD values (red and black, respectively; 20% of the flies in each group; computed for each genetic line as in Fig. 2E). Inset: distribution of MAD scores over the sample.

**Figure S12.**
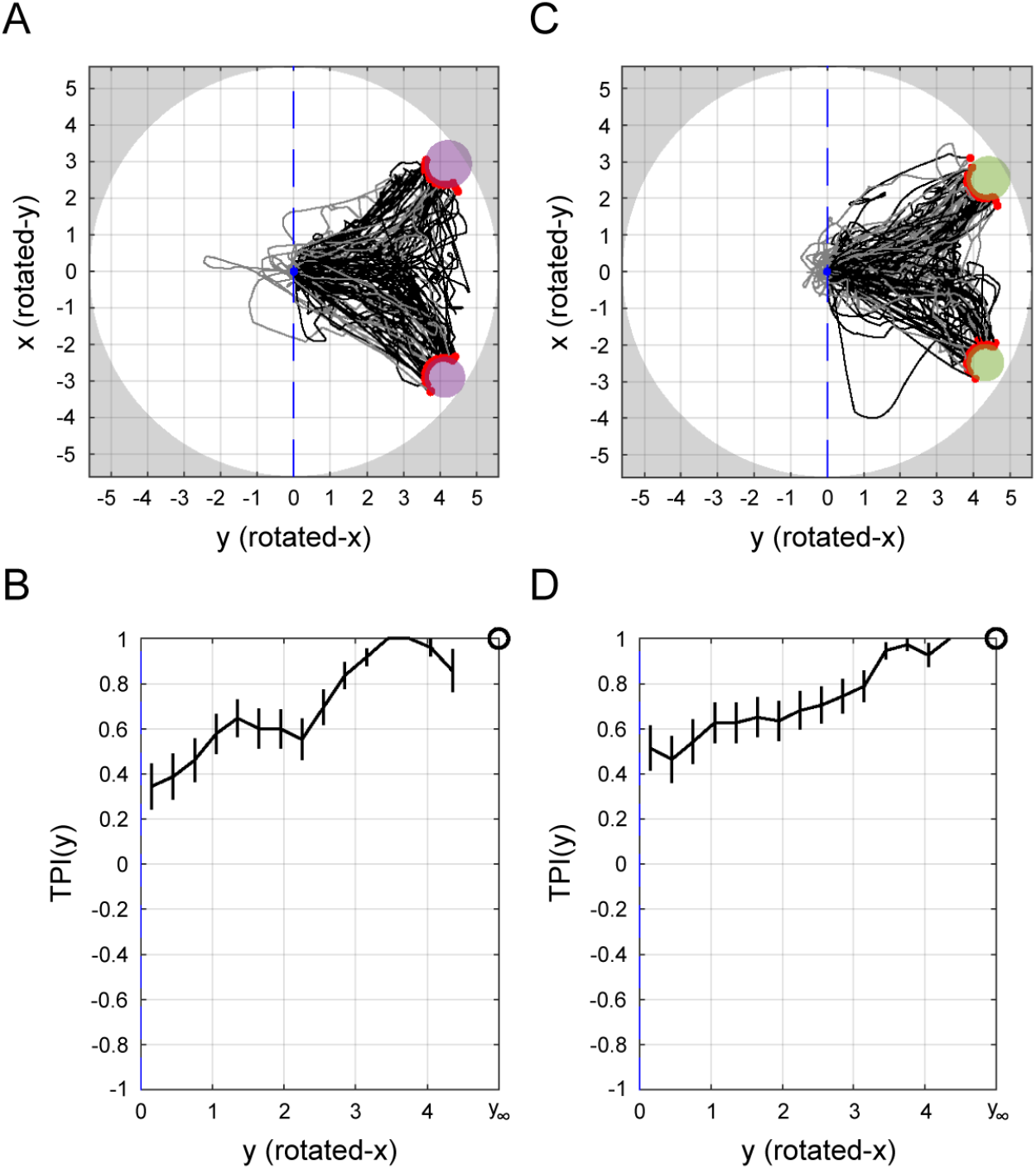
TPI of flies (*n* = 30) for the decision task in the study of Sridhar and colleagues (2021). Flies navigated a circular arena towards identical, equidistant targets. (A) Trajectories of all flies for targets (purple) that were approximately 70 degrees from the initial position of the fly (*n*_*trials*_= 111). Black: trajectories with *min*(*y*) >0 (*n*_*trials*_= 85). Gray: trajectories with min(y) < 0 (*n*_*trials*_= 26). (B) TPI, computed over all the black trajectories in A (*n*_*trials*_= 111). Error bars denote SE. (C) As in A, for targets (green) that were approximately 60 degrees from the initial position of the fly (*n*_*trials*_= 111; black: *n*_*trials*_= 74, gray: *n*_*trials*_= 37). (D) As in B, for the black trajectories in C (*n*_*trials*_= 74). Note that due to the small number of trials per fly, the TPI curves in B and D are computed over all trials in a condition.

**Figure S13.**
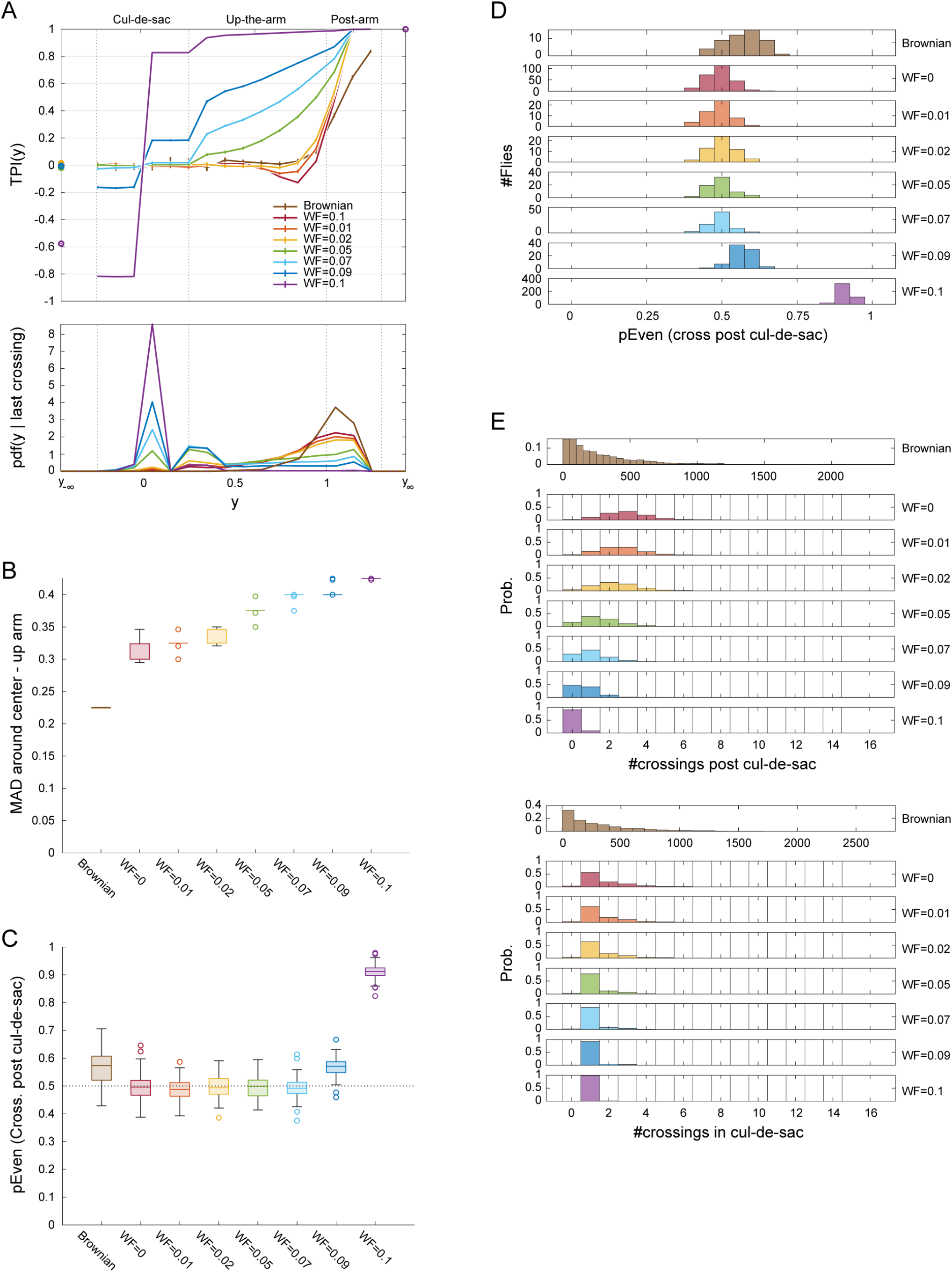
Agent based modeling (ABM). (A) Average TPI curves in the spatial domain and corresponding PDFs of last midline-crossings across AMBs: Brownian ABM (brown, *n* = 50), *WF* =0 (red, *n* = 251), *WF* =0.01 (orange, *n* = 51), *WF* =0.02 (yellow, *n* = 54), *WF* =0.05 (green, *n* = 66), *WF* =0.07 (cyan, *n* = 72), *WF* =0.09 (blue, *n* = 78), and *WF* =0.1 (purple, *n* = 457). Top: Average TPI curve in the spatial domain of each ABM (note the narrow abscisa compared to Fig. 4A, top). Error bars denote SEM. Bottom: PDF of last midline-crossing (LMC), *PDF*(*y* |*LMC*), across all trials and flies in each ABM. (B) Global lateral tendencies. Box plot depicting *MAD*(*x*|0.34 < *y* < 1) scores (computed as in Fig. 4B, top) for each sample. For each box, the central mark indicates the median, bottom and top edges of the box indicate the 25th and 75th percentiles, whiskers extend to the most extreme data points not considered outliers, and outliers are plotted individually using symbols. (C) Local lateral tendencies. Box plot (as in B, bottom) depicting *P*_*Even*_ across each sample. (D) Distributions of *P*_*Even*_, the fraction of trials with even #*crossings* across agents in each ABM. (E) Distributions of the number of midline-crossings post cul-de-sac (left, #*crossings*) and within the cul-de-sac (right) across all trials made by agents in a given ABM (*n*_*trials*_ is 3656, 37458, 7840, 8426, 11957, 14185, 15680, 88564 for the Brownian ABM, *WF* =0, *WF* =0.01, *WF* =0.02, *WF* =0.05, *WF* =0.07, *WF* =0.09 and *WF* =0.1, respectively).

**Figure S14.**
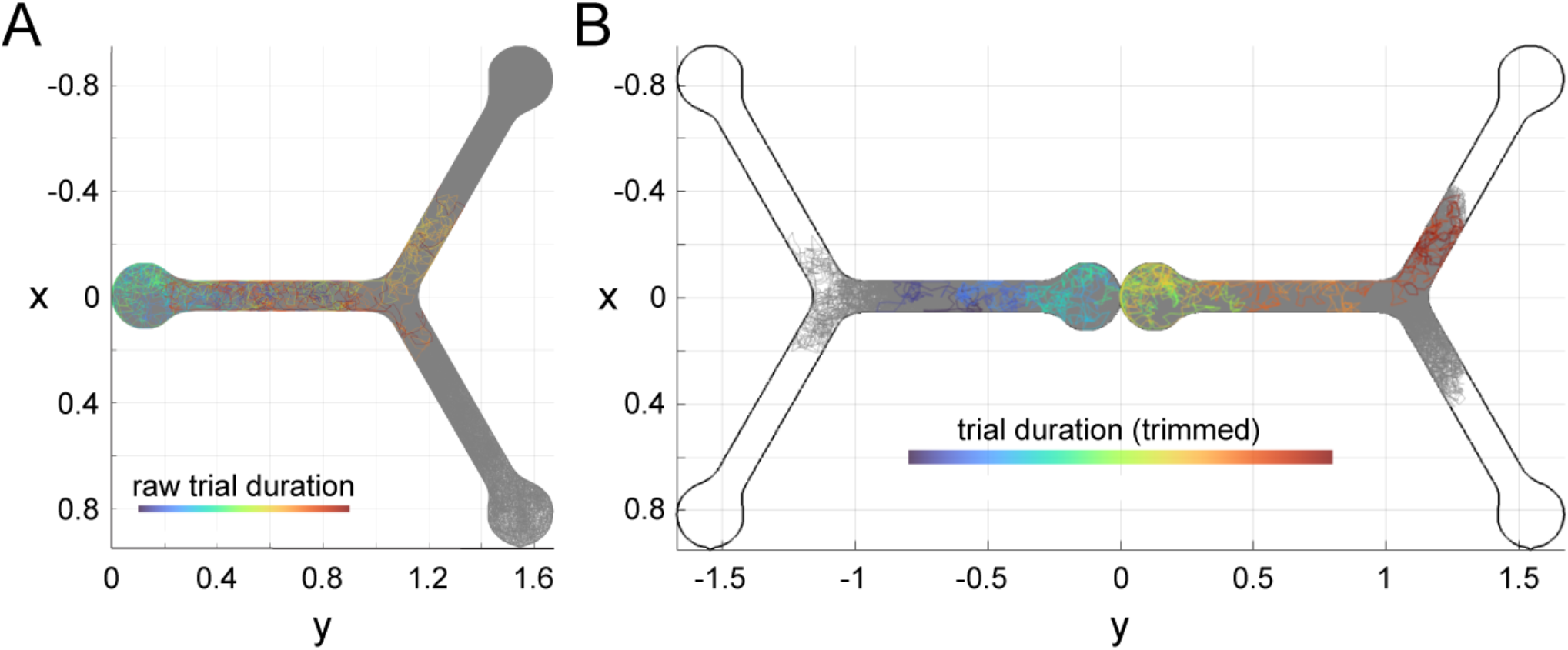
Trajectory trimming and correction for the Brownian ABM simulations. (A) Trajectory of an example trial for one simulated fly (color temperature denotes time progression), overlayed over multiple trajectories of its remaining bottom trials (gray). (B) Corrected and trimmed trajectory. The trajectory in (A) is trimmed such that it includes motion from the first arrival (from above) at the vertical center (*y*∼0.65) of the bottom arm, until the first time *y* = 1.3 is reached from below. The trimmed trajectory is then corrected (as in Fig. 1B-C).

**Figure S15.**
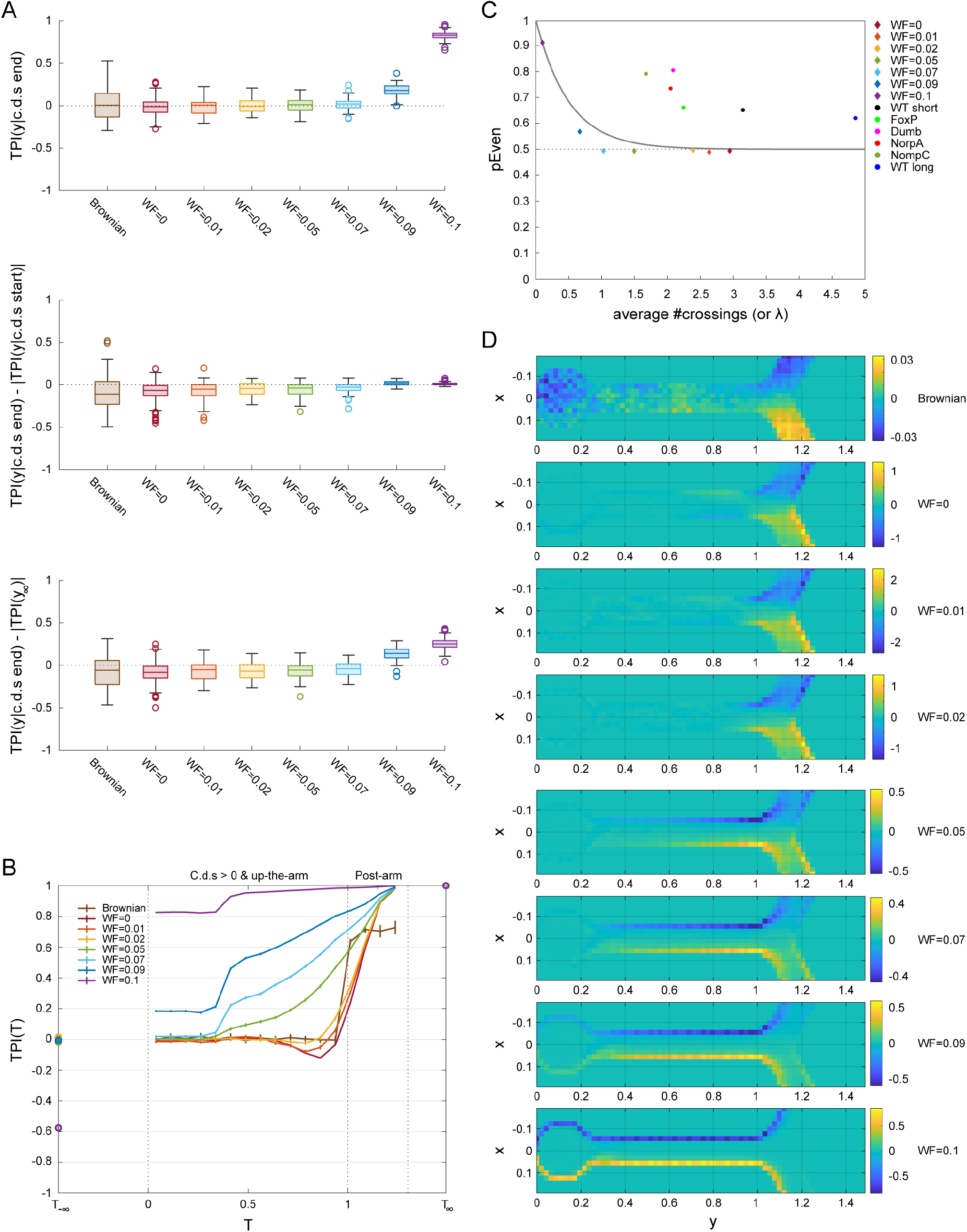
Agent based modeling (ABM). (A) TPI increase in the cul-de-sac across ABMs: Brownian ABM (brown, *n* = 50), *WF* =0 (red, *n* = 251), *WF* =0.01 (orange, *n* = 51), *WF* =0.02 (yellow, *n* = 54), *WF* =0.05 (green, *n* = 66), *WF* =0.07 (cyan, *n* = 72), *WF* =0.09 (blue, *n* = 78), and *WF* =0.1 (purple, *n* = 457). Computed as in S6A. (B) Average TPI curves in the temporal domain across ABMs. Computed as in S8. Note the narrow abscisa compared to Fig. S8. (C) Poisson predictions *vs* observed parity tendencies. Symbols: observed *P*_*Even*_ as a function of the observed average #*crossings* across all trials in each of the genetic lines (diamonds) or ABMs (circles). Black curve: Poisson prediction. Expected *P*_*Even*_(*λ*) under the assumption that #*crossings* in a trial is Poisson distributed with *λ* (the expected value). Genetic lines’ *n*_*trials*_ and ABMs’ colors: 7904 black, 34764 green, 12391 magrenta, 54711 red, 8255 purple, and 7360 blue for WT, FoxP, dumb, NorpA, NompC, and WT in the long maze, respectively. ABM *n*_*trials*_ and colors: 3656 brown, 37458 red, 7840 orange, 8426 yellow, 11957 green, 14185 cyan, 15680 blue, and 88564 purple for the Brownian ABM, *WF* =0, *WF* =0.01, *WF* =0.02, *WF* =0.05, *WF* =0.07, *WF* =0.09 and *WF* =0.1, respectively. (D) Heat maps (as in Fig. 2D, bottom) depicting the difference between the bivariate histograms of right and left turn-decisions for upward traversal (*y* >0). Color temperature denotes probability (colorbar).

## Notes

https://doi.org/10.5281/zenodo.13352340

### Competing Interest Statement

The authors have declared no competing interest.

## References

1. Hanks, T. D., Kopec, C. D., Brunton, B. W., Duan, C. A., Erlich, J. C., & Brody, C. D. (2015). Distinct relationships of parietal and prefrontal cortices to evidence accumulation. Nature 2015 520:7546, 520(7546), 220–223. 10.1038/NATURE14066

2. Giang, T., He, J., Belaidi, S., & Scholz, H. (2017). Key odorants regulate food attraction in Drosophila melanogaster. Frontiers in Behavioral Neuroscience, 11. 10.3389/FNBEH.2017.00160

3. Gibson, W. T., Gonzalez, C. R., Fernandez, C., Ramasamy, L., Tabachnik, T., Du, R. R., Felsen, P. D., Maire, M. R., Perona, P., & Anderson, D. J. (2015). Behavioral responses to a repetitive visual threat stimulus express a persistent state of defensive arousal in drosophila. Current Biology, 25(11), 1401–1415. 10.1016/j.cub.2015.03.058

4. Kempraj, V., Park, S. J., & Taylor, P. W. (2020). Forewarned is forearmed: Queensland fruit flies detect olfactory cues from predators and respond with predator-specific behaviour. Scientific Reports, 10(1), 1–9. 10.1038/s41598-020-64138-6

5. Selen, L. P. J., Shadlen, M. N., & Wolpert, D. M. (2012). Deliberation in the Motor System: Reflex Gains Track Evolving Evidence Leading to a Decision. The Journal of Neuroscience, 32(7), 2276. 10.1523/JNEUROSCI.5273-11.2012

6. Vinauger, C., van Breugel, F., Locke, L. T., Tobin, K. K. S., Dickinson, M. H., Fairhall, A. L., Akbari, O. S., & Riffell, J. A. (2019). Visual-Olfactory Integration in the Human Disease Vector Mosquito Aedes aegypti. Current Biology, 29(15), 2509-2516.e5. 10.1016/j.cub.2019.06.043

7. Lebovich, L., Yunerman, M., Scaiewicz, V., Loewenstein, Y., & Rokni, D. (2021). Paradoxical relationship between speed and accuracy in olfactory figure-background segregation. PLOS Computational Biology, 17(12), e1009674. 10.1371/JOURNAL.PCBI.1009674

8. Bahl, A., & Engert, F. (2020). Neural circuits for evidence accumulation and decision making in larval zebrafish. Nature Neuroscience, 23(1), 94–102. 10.1038/s41593-019-0534-9

9. Huk, A. C., & Shadlen, M. N. (2005). Neural Activity in Macaque Parietal Cortex Reflects Temporal Integration of Visual Motion Signals during Perceptual Decision Making. Journal of Neuroscience, 25(45), 10420–10436. 10.1523/JNEUROSCI.4684-04.2005

10. Shadlen, M. N., & Kiani, R. (2013). Decision making as a window on cognition. Neuron, 80(3), 791–806. 10.1016/J.NEURON.2013.10.047

11. Stüttgen, M. C., & Schwarz, C. (2010). Integration of Vibrotactile Signals for Whisker-Related Perception in Rats Is Governed by Short Time Constants: Comparison of Neurometric and Psychometric Detection Performance. Journal of Neuroscience, 30(6), 2060–2069. 10.1523/JNEUROSCI.3943-09.2010

12. Jankowski, M. M., Polterovich, A., Kazakov, A., Niediek, J., & Nelken, I. (2023). An automated, low-latency environment for studying the neural basis of behavior in freely moving rats. BMC Biology, 21, 172. 10.1186/s12915-023-01660-9

13. Lin, S., Senapati, B., & Tsao, C. H. (2019). Neural basis of hunger-driven behaviour in Drosophila. Open Biology, 9(3). 10.1098/RSOB.180259

14. Liu, H., Yang, W., Wu, T., Duan, F., Soucy, E., Jin, X., & Zhang, Y. (2018). Cholinergic Sensorimotor Integration Regulates Olfactory Steering. Neuron, 97(2), 390-405.e3. 10.1016/j.neuron.2017.12.003

15. Kim, J., Kang, H., Lee, Y.-B., Lee, B., & Lee, D. (2023). A quantitative analysis of spontaneous alternation behaviors on a Y-maze reveals adverse effects of acute social isolation on spatial working memory. Scientific Reports, 13(1), 14722. 10.1038/s41598-023-41996-4

16. Werkhoven, Z., Bravin, A., Skutt-Kakaria, K., Reimers, P., Pallares, L. F., Ayroles, J., & de Bivort, B. L. (2021). The structure of behavioral variation within a genotype. ELife, 10. 10.7554/eLife.64988

17. Sridhar, V. H., Li, L., Gorbonos, D., Nagy, M., Schell, B. R., Sorochkin, T., Gov, N. S., & Couzin, I. D. (2021). The geometry of decision-making in individuals and collectives. Proceedings of the National Academy of Sciences, 118(50), e2102157118. 10.1073/pnas.2102157118

18. Wijnen, K., Genzel, L., & van der Meij, J. (2024). Rodent maze studies: from following simple rules to complex map learning. Brain Structure and Function 2024 229:4, 229(4), 823–841. 10.1007/S00429-024-02771-X

19. Pull, C. D., Petkova, I., Watrobska, C., Pasquier, G., Perez Fernandez, M., & Leadbeater, E. (2022). Ecology dictates the value of memory for foraging bees. Current Biology : CB, 32(19), 4279-4285.e4. 10.1016/J.CUB.2022.07.062

20. Dupret, D., O’Neill, J., Pleydell-Bouverie, B., & Csicsvari, J. (2010). The reorganization and reactivation of hippocampal maps predict spatial memory performance. Nature Neuroscience 2010 13:8, 13(8), 995–1002. 10.1038/nn.2599

21. Alisch, T., Crall, J. D., Kao, A. B., Zucker, D., & de Bivort, B. (2018). MAPLE (Modular automated platform for large-scale experiments), a robot for integrated organism-handling & phenotyping. ELife, 7. 10.7554/ELIFE.37166

22. Ohyama, T., Schneider-Mizell, C. M., Fetter, R. D., Aleman, J. V., Franconville, R., Rivera-Alba, M., Mensh, B. D., Branson, K. M., Simpson, J. H., Truman, J. W., Cardona, A., & Zlatic, M. (2015). A multilevel multimodal circuit enhances action selection in Drosophila. Nature 2015 520:7549, 520(7549), 633–639. 10.1038/nature14297

23. Liu, P., Chen, B., & Wang, Z. W. (2020). GABAergic motor neurons bias locomotor decision-making in C. elegans. Nature Communications 2020 11:1, 11(1), 1–19. 10.1038/s41467-020-18893-9

24. Seibenhener, M. L., & Wooten, M. C. (2015). Use of the Open Field Maze to Measure Locomotor and Anxiety-like Behavior in Mice. Journal of Visualized Experiments : JoVE, 96, 52434. 10.3791/52434

25. Mai, B., Sommer, S., & Hauber, W. (2012). Motivational states influence effort-based decision making in rats: The role of dopamine in the nucleus accumbens. Cognitive, Affective and Behavioral Neuroscience, 12(1), 74–84. 10.3758/S13415-011-0068-4

26. Thomas, E., Snyder, P. J., Pietrzak, R. H., & Maruff, P. (2014). Behavior at the Choice Point: Decision Making in Hidden Pathway Maze Learning. Neuropsychology Review, 24(4), 514–536. 10.1007/s11065-014-9272-7

27. Cleal, M., Fontana, B. D., Ranson, D. C., McBride, S. D., Swinny, J. D., Redhead, E. S., & Parker, M. O. (2020). The Free-movement pattern Y-maze: A cross-species measure of working memory and executive function. Behavior Research Methods 2020 53:2, 53(2), 536–557. 10.3758/S13428-020-01452-X

28. Warden, C. J. (1929). A Standard Unit Animal Maze for General Laboratory Use. Pedagogical Seminary and Journal of Genetic Psychology, 36(1), 174–176.

29. Dylda, E., & Wang, K. H. (2022). Prior actions influence cost–benefit-related decision-making during mouse foraging behaviours. European Journal of Neuroscience, 56(2), 3861–3874. 10.1111/EJN.15689

30. Lacombrade, M., Doblas-Bajo, M., Rocher, N., Tourrain, Z., Navarro, E., Lubat, C., Vogelweith, F., Thiéry, D., & Lihoreau, M. (2023). Flexible visual learning in nectar-foraging hornets. Behavioral Ecology and Sociobiology, 77(7), 1–7. 10.1007/S00265-023-03349-Z

31. Woodley, C. M., Urbanczyk, A. C., Smith, D. L., & Lemasson, B. H. (2019). Integrating Visual Psychophysical Assays within a Y-Maze to Isolate the Role that Visual Features Play in Navigational Decisions. Journal of Visualized Experiments, 2019(147). 10.3791/59281

32. Ayroles, J. F., Buchanan, S. M., O’Leary, C., Skutt-Kakaria, K., Grenier, J. K., Clark, A. G., Hartl, D. L., & de Bivort, B. L. (2015). Behavioral idiosyncrasy reveals genetic control of phenotypic variability. Proceedings of the National Academy of Sciences, 112(21), 6706–6711. 10.1073/pnas.1503830112

33. Buchanan, S. M., Kain, J. S., & de Bivort, B. L. (2015). Neuronal control of locomotor handedness in Drosophila. Proceedings of the National Academy of Sciences, 112(21), 6700– 6705. 10.1073/pnas.1500804112

34. Adel, M., & Griffith, L. C. (2021). The Role of Dopamine in Associative Learning in Drosophila: An Updated Unified Model. Neuroscience Bulletin, 37(6), 831–852. 10.1007/S12264-021-00665-0

35. Lesar, A., Tahir, J., Wolk, J., & Gershow, M. (2021). Switch-like and persistent memory formation in individual Drosophila larvae. ELife, 10. 10.7554/eLife.70317

36. Yeshurun, Y., Carrasco, M., & Maloney, L. T. (2008). Bias and sensitivity in two-interval forced choice procedures: Tests of the difference model. Vision Research, 48(17), 1837–1851. 10.1016/J.VISRES.2008.05.008

37. Ashourian, P., & Loewenstein, Y. (2011). Bayesian Inference Underlies the Contraction Bias in Delayed Comparison Tasks. PLOS ONE, 6(5), e19551. 10.1371/JOURNAL.PONE.0019551

38. Raviv, O., Ahissar, M., & Loewenstein, Y. (2012). How Recent History Affects Perception: The Normative Approach and Its Heuristic Approximation. PLOS Computational Biology, 8(10), e1002731. 10.1371/JOURNAL.PCBI.1002731

39. Urai, A. E., de Gee, J. W., Tsetsos, K., & Donner, T. H. (2019). Choice history biases subsequent evidence accumulation. ELife, 8. 10.7554/ELIFE.46331

40. Thorndike, E. L. (1911). Animal intelligence; experimental studies. In Animal intelligence; experimental studies. The Macmillan Company. 10.5962/bhl.title.55072

41. Skinner, B. F. (1938). The behavior of organisms: an experimental analysis. Appleton-Century. https://psycnet.apa.org/record/1939-00056-000

42. Ferster, C. B., & Skinner, B. F. (1957). Schedules of reinforcement. In Schedules of reinforcement. Appleton-Century-Crofts. 10.1037/10627-000

43. Shteingart, H., & Loewenstein, Y. (2014). Reinforcement learning and human behavior. Current Opinion in Neurobiology, 25, 93–98. 10.1016/J.CONB.2013.12.004

44. Mongillo, G., Shteingart, H., & Loewenstein, Y. (2014). The misbehavior of reinforcement learning. Proceedings of the IEEE, 102(4), 528–541. 10.1109/JPROC.2014.2307022

45. Kain, J. S., Stokes, C., & de Bivort, B. L. (2012). Phototactic personality in fruit flies and its suppression by serotonin and white. Proceedings of the National Academy of Sciences of the United States of America, 109(48), 19834–19839. 10.1073/pnas.1211988109

46. Gorostiza, E. A., Colomb, J., & Brembs, B. (2016). A decision underlies phototaxis in an insect. Open Biology, 6(12), 160229. 10.1098/rsob.160229

47. de Bivort, B., Buchanan, S., Skutt-Kakaria, K., Gajda, E., O’leary, C., Reimers, P., Akhund-Zade, J., Senft, R., Maloney, R., Ho, S., Werkhoven, Z., & Smith, M. A.-Y. (2019). Precise quantification of behavioral individuality from 80 million decisions across 183,000 flies. 10.1101/2021.12.15.472856

48. Lebovich, L., Darshan, R., Lavi, Y., Hansel, D., & Loewenstein, Y. (2019). Idiosyncratic choice bias naturally emerges from intrinsic stochasticity in neuronal dynamics. Nature Human Behaviour, 3(11), 1190–1202. 10.1038/s41562-019-0682-7

49. Churgin, M. A., Lavrentovich, D. O., Smith, M. A., Gao, R., Boyden, E. S., & Bivort, B.de. (2023). Neural correlates of individual odor preference in Drosophila. ELife, 12. 10.7554/ELIFE.90511.1

50. DasGupta, S., Ferreira, C. H., & Miesenböck, G. (2014). FoxP influences the speed and accuracy of a perceptual decision in Drosophila. Science, 344(6186), 901–904. 10.1126/science.1252114

51. Groschner, L. N., Chan Wah Hak, L., Bogacz, R., DasGupta, S., & Miesenböck, G. (2018). Dendritic Integration of Sensory Evidence in Perceptual Decision-Making. Cell, 173(4), 894-905.e13. 10.1016/j.cell.2018.03.075

52. Gold, J. I., & Shadlen, M. N. (2007). The neural basis of decision making. Annual Review of Neuroscience, 30, 535–574. 10.1146/ANNUREV.NEURO.29.051605.113038

53. Ratcliff, R., & McKoon, G. (2008). The Diffusion Decision Model: Theory and Data for Two-Choice Decision Tasks. Neural Computation, 20(4), 873–922. 10.1162/neco.2008.12-06-420

54. Pinto, L., Koay, S. A., Engelhard, B., Yoon, A. M., Deverett, B., Thiberge, S. Y., Witten, I. B., Tank, D. W., & Brody, C. D. (2018). An Accumulation-of-Evidence Task Using Visual Pulses for Mice Navigating in Virtual Reality. Frontiers in Behavioral Neuroscience, 12, 339976. 10.3389/fnbeh.2018.00036

55. Kane, G. A., Senne, R. A., & Scott, B. B. (2023). Rat movements reflect internal decision dynamics in an evidence accumulation task. BioRxiv, 2023.09.11.556575. 10.1101/2023.09.11.556575

56. Molano-Mazón, M., Garcia-Duran, A., Pastor-Ciurana, J., Hernández-Navarro, L., Bektic, L., Lombardo, D., Rocha, J. de la, & Hyafil, A. (2024). Rapid, systematic updating of movement by accumulated decision evidence. BioRxiv. 10.1101/2023.11.09.566389

57. Bett, D., Allison, E., Murdoch, L., Kaefer, K., Wood, E. R., & Dudchenko, P. A. (2012). The neural substrates of deliberative decision making: Contrasting effects of hippocampus lesions on performance and vicarious trial-and-error behavior in a spatial memory task and a visual discrimination task. Frontiers in Behavioral Neuroscience, 6(OCTOBER 2012), 1–32. 10.3389/FNBEH.2012.00070

58. Fontana, B. D., Cleal, M., Clay, J. M., & Parker, M. O. (2019). Zebrafish (Danio rerio) behavioral laterality predicts increased short-term avoidance memory but not stress-reactivity responses. Animal Cognition, 22(6), 1051–1061. 10.1007/S10071-019-01296-9

59. Cleal, M., Fontana, B. D., Double, M., Mezabrovschi, R., Parcell, L., Redhead, E., & Parker, M. O. (2021). Dopaminergic modulation of working memory and cognitive flexibility in a zebrafish model of aging-related cognitive decline. Neurobiology of Aging, 102, 1–16. 10.1016/j.neurobiolaging.2021.02.005

60. Redhead, E. S., Wildschut, T., Oliver, A., Parker, M. O., Wood, A. P., & Sedikides, C. (2023). Nostalgia enhances route learning in a virtual environment. Cognition and Emotion, 37(4), 617– 632. 10.1080/02699931.2023.2185877

61. Gorbonos, D., Gov, N. S., & Couzin, I. D. (2024). Geometrical Structure of Bifurcations during Spatial Decision-Making. PRX Life, 2(1), 013008. 10.1103/PRXLIFE.2.013008

62. Soibam, B., Mann, M., Liu, L., Tran, J., Lobaina, M., Kang, Y. Y., Gunaratne, G. H., Pletcher, S., & Roman, G. (2012). Open-field arena boundary is a primary object of exploration for Drosophila. Brain and Behavior, 2(2), 97. 10.1002/BRB3.36

63. Yan, Z., Zhang, W., He, Y., Gorczyca, D., Xiang, Y., Cheng, L. E., Meltzer, S., Jan, L. Y., & Jan, Y. N. (2012). Drosophila NOMPC is a mechanotransduction channel subunit for gentle-touch sensation. Nature 2012 493:7431, 493(7431), 221–225. 10.1038/NATURE11685

64. Bloomquist, B. T., Shortridge, R. D., Schneuwly, S., Perdew, M., Montell, C., Steller, H., Rubin, G., & Pak, W. L. (1988). Isolation of a putative phospholipase c gene of drosophila, norpA, and its role in phototransduction. Cell, 54(5), 723–733. 10.1016/S0092-8674(88)80017-5

65. Kim, Y. C., Lee, H. G., Seong, C. S., & Han, K. A. (2003). Expression of a D1 dopamine receptor dDA1/DmDOP1 in the central nervous system of Drosophila melanogaster. Gene Expression Patterns, 3(2), 237–245. 10.1016/S1567-133X(02)00098-4

66. Weigel, D., Jürgens, G., Küttner, F., Seifert, E., & Jäckle, H. (1989). The homeotic gene fork head encodes a nuclear protein and is expressed in the terminal regions of the Drosophila embryo. Cell, 57(4), 645–658. 10.1016/0092-8674(89)90133-5

67. Cheng, L. E., Song, W., Looger, L. L., Jan, L. Y., & Jan, Y. N. (2010). The Role of the TRP Channel NompC in Drosophila Larval and Adult Locomotion. Neuron, 67(3), 373–380. 10.1016/J.NEURON.2010.07.004

68. Pak, W. L., Grossfield, J., & Arnold, K. S. (1970). Mutants of the Visual Pathway of Drosophila melanogaster. Nature 1970 227:5257, 227(5257), 518–520. 10.1038/227518B0

69. Kim, Y. C., Lee, H. G., & Han, K. A. (2007). D1 Dopamine Receptor dDA1 Is Required in the Mushroom Body Neurons for Aversive and Appetitive Learning in Drosophila. The Journal of Neuroscience, 27(29), 7640. 10.1523/JNEUROSCI.1167-07.2007

70. Kottler, B., Faville, R., Bridi, J. C., & Hirth, F. (2019). Inverse Control of Turning Behavior by Dopamine D1 Receptor Signaling in Columnar and Ring Neurons of the Central Complex in Drosophila. Current Biology, 29(4), 567. 10.1016/J.CUB.2019.01.017

71. Palazzo, O., Rass, M., & Brembs, B. (2020). Identification of FoxP circuits involved in locomotion and object fixation in Drosophila: FoxP in locomotion and object fixation. Open Biology, 10(12). 10.1098/rsob.200295

72. Neuser, K., Triphan, T., Mronz, M., Poeck, B., & Strauss, R. (2008). Analysis of a spatial orientation memory in Drosophila. Nature 2008 453:7199, 453(7199), 1244–1247. 10.1038/nature07003

73. Ofstad, T. A., Zuker, C. S., & Reiser, M. B. (2011). Visual place learning in Drosophila melanogaster. Nature, 474(7350), 204–209. 10.1038/NATURE10131

74. Kim, I. S., & Dickinson, M. H. (2017). Idiothetic Path Integration in the Fruit Fly Drosophila melanogaster. Current Biology, 27(15), 2227-2238.e3. 10.1016/j.cub.2017.06.026

75. Cheong, H., Siwanowicz, I., & Card, G. M. (2020). Multi-regional circuits underlying visually guided decision-making in Drosophila. Current Opinion in Neurobiology, 65, 77–87. 10.1016/J.CONB.2020.10.010

76. Zhang, K., Guo, J. Z., Peng, Y., Xi, W., & Guo, A. (2007). Dopamine-mushroom body circuit regulates saliency-based decision-making in Drosophila. Science, 316(5833), 1901–1904. 10.1126/science.1137357

77. Fisher, Y. E., Marquis, M., D’Alessandro, I., & Wilson, R. I. (2022). Dopamine promotes head direction plasticity during orienting movements. Nature 2022 612:7939, 612(7939), 316–322. 10.1038/s41586-022-05485-4

78. Mohandasan, R., Iqbal, F. M., Thakare, M., Sridharan, M., & Das, G. (2020). Enhanced odour-associated memory performance with a Y-maze assembly in Drosophila. BioRxiv, 2020.11.18.386128. 10.1101/2020.11.18.386128

79. Mendoza, E., Colomb, J., Rybak, J., Pflüger, H.-J., Zars, T., Scharff, C., & Brembs, B. (2014). Drosophila FoxP Mutants Are Deficient in Operant Self-Learning. PLoS ONE, 9(6), e100648. 10.1371/journal.pone.0100648

80. Ehweiner, A., Duch, C., & Brembs, B. (2024). Wings of Change: aPKC/FoxP-dependent plasticity in steering motor neurons underlies operant self-learning in Drosophila. F1000Research, 13, 116. 10.12688/f1000research.146347.2

81. Milosavljevic, M., Malmaud, J., Huth, A., Koch, C., & Rangel, A. (2010). The Drift Diffusion Model can account for the accuracy and reaction time of value-based choices under high and low time pressure. Judgment and Decision Making, 5(6), 437–449. 10.2139/ssrn.1901533

82. Buerkle, N., & Jeanne, J. M. (2021). Decision making: An analogue implementation of a drift-diffusion computation in the Drosophila mushroom body. Current Biology, 31(22), R1479–R1482. 10.1016/J.CUB.2021.09.039

83. Oliver, A., Wildschut, T., Parker, M. O., Wood, A. P., & Redhead, E. S. (2023). Induction of spatial anxiety in a virtual navigation environment. Behavior Research Methods, 55(7), 3621– 3628. 10.3758/S13428-022-01979-1

84. Werkhoven, Z., Rohrsen, C., Qin, C., Brembs, B., & de Bivort, B. (2019). MARGO (Massively Automated Real-time GUI for Object-tracking), a platform for high-throughput ethology. PLoS ONE, 14(11). 10.1371/JOURNAL.PONE.0224243

85. Lebovich, L., Alisch, T., Redhead, E. S., Parker, M. O., Loewenstein, Y., Couzin, I. D., & de Bivort, B. L. (2024). Spatiotemporal dynamics of locomotor decisions in Drosophila melanogaster [Data set]. Zenodo. 10.5281/zenodo.13352340

